# Multiple waves of viral invasions in Symbiodiniaceae algal genomes

**DOI:** 10.1101/2022.04.12.488082

**Authors:** L. Felipe Benites, Timothy G. Stephens, Debashish Bhattacharya

**Author notes:** Corresponding author L. Felipe Benites - Department of Biochemistry and Microbiology, Rutgers University, New Brunswick, NJ 08901, USA.

## Abstract

Dinoflagellates from the family Symbiodiniaceae are phototrophic marine protists that engage in symbiosis with diverse hosts. Their large and distinct genomes show pervasive gene duplication and large-scale retroposition events. However, little is known about the role and scale of horizontal gene transfer (HGT) in the genomic evolution of this algal family. In other dinoflagellates, higher levels of HGTs have been observed, linked to major genomic transitions, such as the appearance of a viral acquired nucleoprotein that originated *via* HGT from a large DNA algal virus. Previous work showed Symbiodiniaceae from different hosts being actively infected by several viral groups, such as giant DNA viruses and ssRNA viruses, that may play an important role in coral health. This includes a hypothetical latent viral infection, whereby viruses could persist in the cytoplasm or integrate into the host genome as a provirus. This hypothesis received some experimental support however, the cellular localization of putative latent viruses and their taxonomic affiliation are still unknown. In addition, despite the finding of viral sequences in some genomes of Symbiodiniaceae, viral origin, taxonomic breadth, and metabolic potential have not been explored. To address these questions, we searched for evidence of protein sequences of putative viral origin in 13 Symbiodiniaceae genomes. We found 59 candidate viral-derived HGTs that give rise to 12 phylogenies across 10 genomes. We also describe the taxonomic affiliation of these virus-related sequences, their structure, and genomic context. These results lead us to propose a model to explain the origin and fate of Symbiodiniaceae viral acquisitions.

## 1. Introduction

Dinoflagellates from the family Symbiodiniaceae are phototrophic marine protists that engage in symbiosis with diverse hosts such as foraminifera, ciliates, radiolarians, mollusks, and cnidarians such as anemones and most notably corals (LaJeunesse et al. 2018). Their large and distinct genomes show pervasive gene duplication (Aranda et al. 2016; Nand et al. 2021), expansion of lineage specific gene families with unknown functions (González-Pech et al. 2021), and large-scale retroposition events (Song et al. 2017). However, little is known about the role of horizontal gene transfer (HGT) in Symbiodiniaceae evolution and the scale of gene acquisition events (González-Pech et al. 2021). In *Fugacium kawagutii* and *Brevioloum minutum* (formerly *Symbiodinium* clades F and B), ∼0.7% of their genes (246 and 244 sequences respectively), have putatively originated from prokaryote-derived HGT events (Fan et al. 2020).

Higher levels of HGTs have been observed in other dinoflagellates and appear to be linked to major genomic transitions (Wisecaver et al. 2013; Janouškovec et al. 2017). Remarkably, all sequenced dinoflagellate genomes contain a family of viral acquired nucleoproteins, known as the dinoflagellate/viral nucleoproteins (DVNPs), which have a putative role in chromatin regulation (Irwin et al. 2018). In *B. minutum*, 19 divergent DVNPs have been identified and it has been suggested that they are involved in regulating chromosome structure (Shoguchi et al. 2013). Furthermore, it has been proposed that the DVNP protein family originated in the ancestor of dinoflagellates by HGT from a large DNA algal virus, possibly from the family *Phycodnaviridae* (Gornik et al. 2012). Acquisition of the DVNP gene family coincides with massive genomic expansion in this group and may have conferred immunity to infection by the same group of exogenous viruses (Gornik et al. 2019).

Previous work has found cases of Symbiodiniaceae isolated from corals being actively infected by several viral groups, ranging from giant DNA viruses to ssRNA viruses (A.M.S. Correa et al. 2013; Weynberg et al. 2014; Wood-Charlson et al. 2015; A.M.S. Correa et al. 2016; Levin et al. 2017; Messyasz et al. 2020). Most importantly, the presence and abundance of viral sequences were found to be associated with bleached and diseased corals, suggesting that viruses may play a role in coral health and bleaching events by infecting (subsequently killing or disrupting the function of) Symbiodiniaceae cells in the host (Thurber and A.M.S. Correa 2011). Recently, Grupstra et al. (2022) demonstrated the rapid formation of diverse viral major capsid protein (MCP) “aminotypes” (unique amino acid sequences) from a Symbiodiniaceae-infecting dinoflagellate RNA virus in coral colonies exposed to high temperatures (30.3 °C) that are known to cause coral bleaching. This suggests that the switch from a persistent to productive viral infection was triggered by exposure to high temperatures.

Early observations linking symbiotic dinoflagellates from anemones with viruses, stresses, and coral diseases (Wilson et al. 2001) led to the hypothesis that the algal cells harbored latent proviruses. In this type of infection, a virus persists in the cytoplasm as an episome (an extrachromosomal molecule) or becomes integrated into the host genome, replicating as a (latent) provirus, synchronized with the host cell (Hyman and Abedon 2012). After exposure to stress, the latent provirus virus enters a lytic cycle, lysing the host cell and propagating into other susceptible hosts. This hypothesis was partially corroborated by experiments using multiple Symbiodiniaceae isolates from various cnidarian hosts, showing the production of viral-like particles (VLPs) in thermally and ultraviolet (UV) light stressed algal cells (Wilson et al. 2005; Davy et al. 2006; Lohr et al. 2007; Lawrence et al. 2014; Lawrence et al. 2017; Weynberg et al. 2017; Benites et al. 2018). However, the cellular localization of putative latent viruses (i.e., cytoplasmic or genomic) and their precise taxonomic affiliations is still unknown.

In other algal groups, such as the brown algae (Phaeophyceae), double-stranded DNA viruses from *Phycodnaviridae* can persist as integrated proviruses with fragments of the viral genome, interspersed as repeats and pseudogenes throughout the host genome. The viral genome remains latent until it becomes active in reproductive cells after light and temperature stimulation (Bräutigam et al. 1995: 1; Müller et al. 1998; Delaroque and Boland 2008; McKeown et al. 2017). In *Ectocarpus* (strain Ec 32), the genome of a virus was found to be integrated in the host genome however, the viral genes were not expressed and viral particles were not produced (Cock et al. 2010). This is similar to observations in *Feldmannia,* which has viral fragments in its genome but does not show evidence of viral production (Lee et al. 1998). In the red alga (Rhodophyta) *Chondrus crispus*, multiple copies of a partial RNA viral genome were found to be expressed, including a copy of the capsid gene, but there was no apparent virus production or host symptoms (Rousvoal et al. 2016). In green algae (Chlorophyta), although some of their genomes contain numerous giant “endogenous viral elements’’ (EVEs) with spliceosomal introns and segmental gene duplications (Moniruzzaman et al. 2020), the only evidence thus far of viral latency is from the chlorophyte *Cylindrocapsa geminella*, with production of VLPs and lytic infection observed after heat-shock treatment (Hoffman and Stanker 1976).

Recently, Irwin et al. (2022) reported that at least 23 genes in Symbiodiniaceae (Fkaw, Sgor and Smic) show strong support for HGTs with viruses such as Nucleocytoviricota giant viruses (although their analysis did not focus on Symbiodiniaceae), with possible roles in catalysis of ATP-dependent conversion of ribonucleotides, HNH endonuclease, among others. In addition, despite the fact that viral sequences account for < 3% of the *Symbiodinium microadriaticum* genome (Aranda et al. 2016) and ∼0.1% of the *Cladocopium goreaui* genome (Liu et al. 2018), their origin, taxonomic breath, and metabolic potential has not been explored in depth and there has been no previous evidence of viral genome integration into Symbiodiniaceae nuclear DNA. The latter finding would strongly support the proviral hypothesis (Wilson et al. 2001).

Therefore, to address questions about the occurrence of integrated viral sequences in Symbiodiniaceae genomes, and to uncover their possible origins and functional profiles, we searched predicted proteins from the 13 available Symbiodiniaceae genomes (Shoguchi et al. 2013; Lin et al. 2015; Liu et al. 2018; Shoguchi et al. 2018; Chen et al. 2020; González-Pech et al. 2021) for sequences of putative viral origin. We identified 59 candidate viral HGT sequences that formed 12 phylogenies, across 10 genomes. We describe for each of these sequences their taxonomic affiliation with viruses and the composition, structure, and genomic context of their associated genes. Moreover, we searched for corroborative evidence that would support these sequences being HGT-derived and for the presence of viral genome integrations.

## 2. Methods

### 2.1 Screening for viral HGT events in Symbiodiniaceae protein datasets

Each predicted protein from the 13 Symbiodiniaceae genomes (445,306 total) was searched against the RefSeq viral database using DIAMOND (v0.9.22; BLASTp, sensitive mode) (Buchfink et al. 2015) with an *e*-value cutoff of < 1e^-5^, retaining just the best viral hits per query sequence. To exclude false positive hits, we perform another DIAMOND (BLASTp) search against the complete Genbank non-redundant database, using only the Symbiodiniaceae protein queries with hits to the RefSeq viral database. For each query, the top 400 hits were retained and filtered using an *e*-value of < 1e^-5^. For each hit, the full taxonomy of the subject sequence was retrieved for the top 50 filtered hits using a custom perl script (available at: https://github.com/LFelipe-B/Symbiodiniaceae_vHGT_scripts) which uses the subject accession number to fetch the whole taxonomic tree from the Genbank taxonomy suit. Query sequences that had at least one of their top 50 hits to a viral sequence were retained for downstream analysis. Genomes are abbreviated as follows: *Brevioloum minutum* (Bmin); *Cladocopium* sp. C92 (CC92); *Cladocopium goreaui* (Cgor); *Fugacium kawagutii* (Fkaw); *Symbiodinium linucheae* CCMP2456 (Slin_CCMP2456); *S. microadriaticum* (Smic); *S. microadriaticum* 04-503SCI.03 (Smic_04-503SCI.03); *S. microadriaticum* CassKB8 (Smic_CassKB8); *S. natans* CCMP2548 (Snat_CCMP2548); *S. necroappetens* CCMP2469 (Snec_CCMP2469); *S. pilosum* CCMP2461 (Spil_CCMP2461); *S. tridacnidorum* (Stri); and *S. tridacnidorum* CCMP2592 (Stri_CCMP2592).

### 2.2 Inferring pre-candidates for viral HGTs

To determine the scale of viral HGT in Symbiodiniaceae genomes we collected sequences that showed significant similarity to viral sequences by calculating four independent metrics using custom python scripts and the full taxonomic annotations retrieved using the subject accession numbers from the DIAMOND outputs. Each hit to the complete Genbank non-redundant database was categorized as “non-target” if the subject sequence was from a virus and as “target” if the subject sequence was from a eukaryote; hits to Symbiodiniaceae sequences were omitted from the subsequent calculations to avoid hits to sequences from the same or similar species already submitted to GenBank from biasing the analysis. The first metric calculated was the Alien index (AI) (Gladyshev et al. 2008; Rancurel et al. 2017), which assesses how many orders of magnitude there is between the *e*-values of the top target and non-target hits; query sequences were retained if non-target sequences scored AI > 30. The second metric calculated was the HGT index (hU) (Boschetti et al. 2012), defined as the difference between the bitscores of the top non-target and target hits; query sequences with a Hu ≥ 30 were retained. The third metric was calculated by counting the total number of target and non-target hits (collapsing repeated taxa) associated with each query; query sequences were retained if the sum of non-target hits was > 50% of the total number of hits. Finally, the fourth metric was calculated by taking the sum of the non-target hit bitscores and comparing it with the sum of the target hit bitscores, retaining query sequences which had higher non-target bitscore totals compared to their target bitscore totals. Query sequences with scores above the cut offs associated with each of the four metrics were considered pre-candidates of viral HGTs (pre-vHGTs).

### 2.3 Orthogroup clustering of pre-vHGTs and Phylogenetic reconstruction

All Symbiodiniaceae proteins were clustered into orthogroups using OrthoFinder v2.3.3 (Emms and Kelly 2019), with groups containing at least one pre-vHGT sequence extracted for further analysis. We downloaded the subject sequences associated with each of the pre-vHGT DIAMOND hits to the RefSeq viral and Genbank non-redundant databases (accession numbers in Supplementary Table S11). The Symbiodiniaceae proteins in each pre-vHGT-containing orthogroup were combined with the downloaded subject sequences and complemented with subject hits downloaded from a BLASTp search (https://blast.ncbi.nlm.nih.gov) to retrieve the maximum number of subject hits associated with the pre-vHGTs for each group. The combined set of proteins associated with each group were aligned using MAFFT v7.305b (L-INS-i algorithm: --localpair --maxiterate 1000) (Katoh and Standley 2013). The resulting alignments were trimmed using TrimAI v1.2 in automated mode (-automated1) (Capella-Gutiérrez et al. 2009). A Maximum Likelihood phylogenetic tree was inferred from each group alignment using IQ-TREE v2.0.3 (Nguyen et al. 2015), with the best evolutionary model selected by ModelFinder (Kalyaanamoorthy et al. 2017) (-m TEST) and 10,000 ultrafast bootstrap replicates calculated using UFBoot (Minh et al. 2013). Phylogenetic trees were midpoint rooted and screened for topologies in which Symbiodiniaceae query sequences (target) were clustered or nested with viral sequences (non-target) in a clade with > 90% ultrafast bootstrap support using PhySortR (Stephens et al. 2016) (clade.exclusivity = 0.95, min.prop.target = 0.7). Phylogenies were further processed using the packages ggtree (Yu et al. 2017), phangorn (Schliep 2011), and phytools (Revell 2012) in the R environment (version 3.4.2). A final manual curation step was implemented, discarding orthogroups with ambiguous phylogenetic signals: cases where there was an over representation of non-viral sequences or excessively long branch lengths were considered as having weak evidence of viral HGT and were discarded. The phylogenetic groups that remained after filtering were considered to contain Symbiodiniaceae sequences (hereinafter referred to as vHGTs) with strong evidence of having originated from viral HGT.

### 2.4 Functional analysis of Symbiodiniaceae vHGTs

To gain insights into the functional landscape of viral acquired sequences, we analyzed protein family membership and gene ontology (GO) terms for the vHGTs by re-annotating individual sequences using the InterProScan online suite (https://www.ebi.ac.uk/interpro/) and QuickGO (https://www.ebi.ac.uk/QuickGO/annotations). In some instances, we also subjected the vHGTs to a BLASTp search against the UniProtKB database (https://www.uniprot.org/uniprot/).

### 2.5 Compositional and structural analysis of Symbiodiniaceae vHGTs

We retrieved information about the GC-content, length of coding sequences (CDSs), presence of introns, intron length, and location along the scaffold of the gene models associated with the vHGTs. For each of these features, the difference between the mean values calculated for the vHGTs was compared against a background of all Symbiodiniaceae genes using the Welch Two Sample t-test in R. The presence of a dinoflagellate spliced leader (DinoSL) sequence upstream of the first exon of the vHGT genes in the genome was assessed to identify if mRNA recycling (Slamovits and Keeling 2008) has played a role in the evolution of these genes. DinoSL and relic DinoSL sequences were identified by an approach similar to that used by (Stephens et al. 2020). Briefly, the DinoSL sequence (CCGTAGCCATTTTGGCTCAAG) and relic DinoSL sequences (which are composed of two or more DinoSL sequences joined together at their canonical splice sites) were searched against each Symbiodiniaceae genome using BLASTn v2.10.1 (-max_target_seqs 1000000 -task blastn-short -evalue 1000). Hits were retained if they started ≤ 5 bp from the 5′-end of the query sequence (which is the position of the conserved canonical splice site) and ≤ 2 bp from the 3′-end of the query. The proximity of the identified DinoSL and relic DinoSL sequences to vHGT genes within the genome was assessed using bedtools v2.29.2 (“bedtools closest -s -D a -id”). Only cases where a DinoSL or relic DinoSL were identified on the same strand as the vHGT gene, were upstream or overlapping with the vHGT, and were < 2 Kbp from the first (most upstream) exon of the vHGT gene were reported.

### 2.6 Gene expression analysis and gene model confirmation of Symbiodiniaceae vHGTs

We retrieved RNA-seq files from the Sequence Read Archive (SRA) database (https://www.ncbi.nlm.nih.gov/sra) (accession list in Supplementary Table S10) to study the expression of vHGTs and to validate the vHGTs gene models. When possible (with the exception of Snec, which lacks RNA-seq data), we used RNA-seq data generated from the same isolate to verify the gene modes in the genome. The RNA-seq data was downloaded from NCBI using Fastq-dump v2.9.6 (https://trace.ncbi.nlm.nih.gov/Traces/sra/sra.cgi?view=toolkit_doc&f=fastq-dump), trimmed using Cutadapt v1.18 (Martin 2011) (quality-cutoff 20 and minimum-length 25), and mapped against the associated reference genome using HISAT2 v2.2.1 (Kim et al. 2019). The resulting alignment files were sorted and indexed using SAMtools v1.11 (Li et al. 2009), before being visualized using the Integrative Genomics Viewer (IGV) v2.12.3 tool (Robinson et al. 2011).

The gene models associated with the putative sites of *Symbiodinium* +ssRNA virus integration into the host genome were further analyzed for internal stop codons and frameshift mutation to determine if these sequences had become pseudogenes or if they are still potentially functional. For each viral +ssRNA gene of interest, the genome sequence, starting 2 kbp upstream and ending 2 kbp downstream, was extracted using seqkit v0.15.0 (-u 2000 -d 2000) (Shen et al. 2016). These sequences were used in alignments that included known RNA-dependent RNA polymerase (YP_009337004.1 and YP_009342067.1; top hits using online BLASTP against the viral gene models) and major capsid (AOG17586.1) proteins using Exonerate v2.3.0 (--model protein2genome --exhaustive yes) (Slater and Birney 2005).

## 3. Results

### 3.1 Screening Symbiodiniaceae sequences using four different sequence similarity HGT scoring metrics

The four HGT metrics (see Methods) were applied to Symbiodiniaceae proteins with hits to viral sequences in RefSeq to produce the list of viral HGT pre-candidates (pre-vHGTs). We identified additional viral HGT candidates by constructing orthogroups and taking all sequences that were in an orthogroup with a pre-vHGT. The Symbiodiniaceae sequences from each group, or individually if single sequences, were combined with the sequences retrieved from the top hits of each pre-vHGT against a taxonomically diverse database and expanded to contain the maximum number of subject hits with BLASTp; for each group of combined sequences a multiple sequence alignment was created that was then used for phylogeny reconstruction. The resulting phylogenies were filtered, retaining only those in which viruses were overrepresented and in which viral sequences were nested with Symbiodiniaceae sequences in a clade that had UFBoot branch support > 90%. Our workflow resulted in the identification of 59 vHGT sequences (that comprise 12 phylogenies) that are spread across the 10 Symbiodiniaceae genomes, ranging from n = 14 sequences in Smic to n = 1 in the Cgor and Slin_CCMP2456 genomes. The majority of vHGTs were identified in the genomes of species from the *Symbiodinium* genus, and were absent from the genomes of CC92, Fkaw and the free living Spil_CCMP2461 species.

### 3.2 Taxonomic distribution of vHGTs sequences in Symbiodiniaceae

The taxonomic distribution of the vHGTs was assessed at all classification levels using the top viral hit associated with each sequence and associated taxonomic information in the NCBI database. The number of viral sequences grouped at the order level in each of the genomes is shown in Fig. 1. The vHGTs were derived from both DNA and RNA viruses, however, the majority were annotated as Unclassified RNA viruses (NCBI Taxonomy ID: 1922348; n = 44 sequences). The genome with the most Unclassified RNA viruses vHGTs was *Symbiodinium microadriaticum* (n = 14), followed by the isolates *S. microadriaticum* KB8 (n = 11), and *S. microadriaticum* 04-503SCI.03 (n = 6). The second most abundant group of vHGTs (n = 11) was similar to giant viruses from the order Pimascovirales (Phylum Nucleotycoviricota; formerly NCLDVs). Finally, the third group of vHGTs (n = 4) was annotated to negative-strand RNA viruses from the Mononegavirales (full annotation in Supplementary Table S2).

**Fig. 1:**
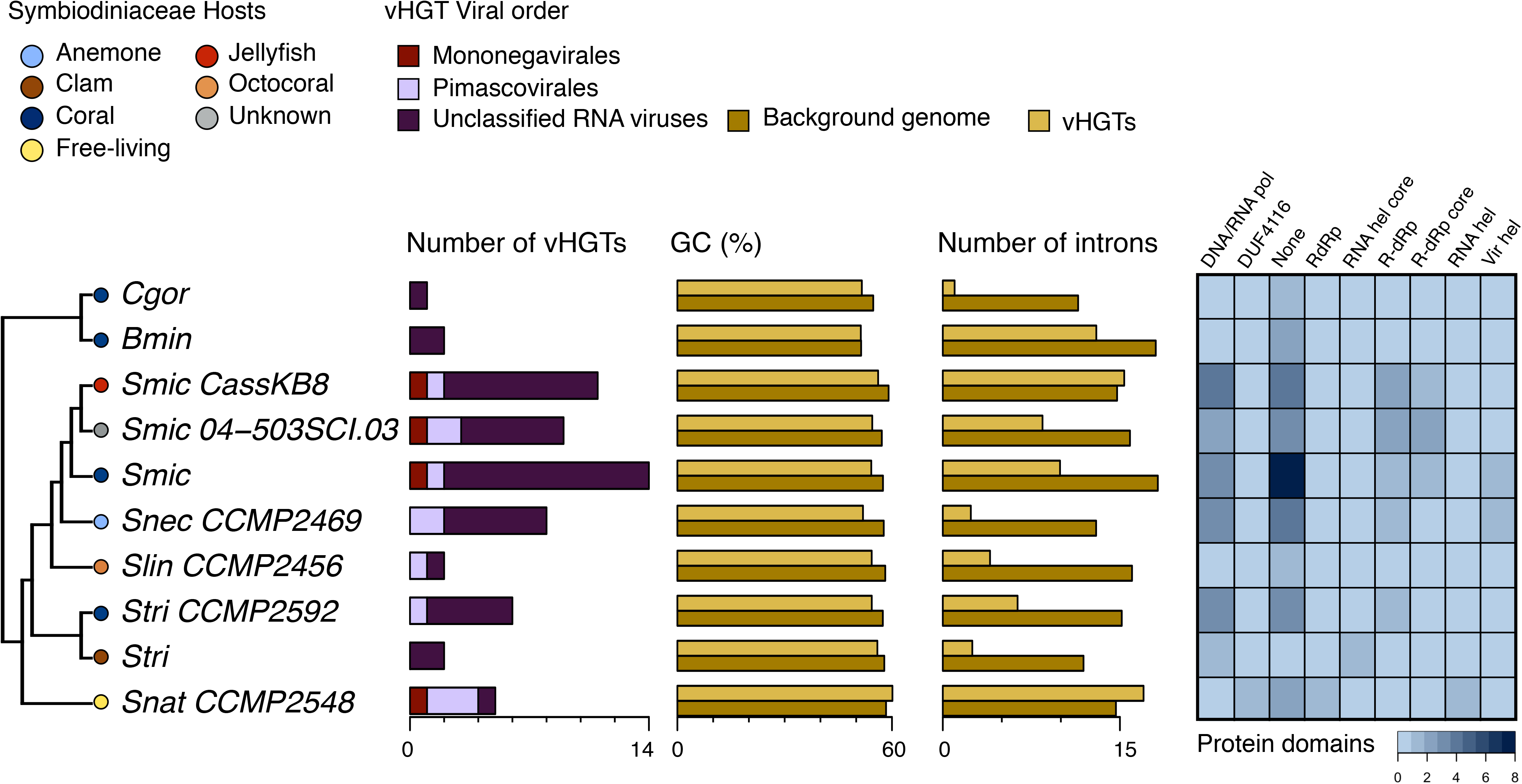
Main characteristics of genes in Symbiodiniaceae genomes that were acquired by viral horizontal gene transfer (vHGT). Cladogram depicting phylogenetic relationship of Symbiodiniaceae genomes in which vHGTs were identified (left); the hosts associated with each Symbiodiniaceae species are represented by colored circles (top left key). The total number of vHGTs and their putative taxonomic origins (at the order level; center key), the coding sequence (CDS) percent Guanine-Cytosine (GC%) content, the average number of introns, and the total number of associated protein domains are shown for the vHGTs identified in each genome. The CDS GC% content and average number of introns is shown separately for the vHGTs and the background Symbiodiniaceae sequences (i.e., all Symbiodiniaceae CDS or genes [respectively] excluding the vHGTs). Abbreviations: *Brevioloum minutum* (Bmin); *Cladocopium* sp. C92 (CC92); *Cladocopium goreaui* (Cgor); *Fugacium kawagutii* (Fkaw); *Symbiodinium linucheae* CCMP2456 (Slin_CCMP2456); *S. microadriaticum* (Smic); S*. microadriaticum* 04-503SCI.03 (Smic_04-503SCI.03); *S. microadriaticum* CassKB8 (Smic_CassKB8); *S. natans* CCMP2548 (Snat_CCMP2548); *S. necroappetens* CCMP2469 (Snec_CCMP2469); *S. pilosum* CCMP2461 (Spil_CCMP2461); *S. tridacnidorum* (Stri); *S. tridacnidorum* CCMP2592 (Stri_CCMP2592); DNA/RNA pol (DNA/RNA polymerase superfamily); DUF4116 (Domain of unknown function DUF4116); None (None predicted); RdRp (RNA dependent RNA polymerase); RNA hel core (RNA virus helicase core domain); R-dRP (RNA-directed RNA polymerase); R-dRp core (RNA-directed RNA polymerase catalytic domain); RNA hel (Viral (Superfamily 1) RNA helicase); Vir hel (Viral helicase1).

### 3.3 Functional annotation and expression of vHGTs sequences in Symbiodiniaceae

We found that the most abundant predicted functional annotation associated with the vHGTs is the DNA/RNA polymerase superfamily (n = 12), followed by RNA-directed RNA polymerase (n = 11), and viral RNA helicase (n =3). Eleven vHGTs had RNA-directed RNA polymerase annotations (GO:0003968), followed by ATP binding (GO:0005524), viral RNA genome replication (GO:0039694), and mRNA capping (GO:0006370) as the most abundant GO terms. There were no protein family membership or gene ontology terms (GO terms) predicted for ten of the vHGTs (Fig. 1) (full annotation in Supplementary Table S2).

We also noted six vHGTs occurring in Smic, Smic_04-503SCI.03, Smic_CassKB8 and Stri_CCMP2592 that have two predicted domains, one of viral origin and the other of eukaryotic origin, suggesting the possibility that they might be chimeric genes (Méheust et al. 2018) (Supplementary Table S9). At the 5’-end of these proteins is a predicted Clavaminate synthase-like or a Winged helix DNA-binding domain and at the 3’-end is a DNA/RNA polymerase domain. Aligned RNA-seq reads were visualized and compared manually against all the predicted vHGT gene models to assess if these sequences are being expressed with at least one read mapping to the gene model. This analysis showed that only 13/59 vHGTs had evidence of read mapping (Supplementary Table S2). This could be explained by the experimental conditions used to generate the RNA-seq data, or alternatively, the vHGTs are non-functional. Because there was also no expression of the putative chimeric domain genes we could not validate if these sequences are true chimeras or artifacts of gene prediction.

### 3.4 Phylogenetic profile of vHGTs sequences in Symbiodiniaceae

#### 3.4.1 Unclassified RNA viruses

The majority of vHGTs were grouped into nine phylogenies (n = 44 sequences); these sequences are all putatively related to unclassified RNA viruses (Shi et al. 2016) (Fig. 2). The taxonomy of top viral hits contained the Wenzhou weivirus-like virus (n = 15), Beihai sobemo-like virus (n = 7), Beihai weivirus-like virus (n = 7), Beihai narna-like virus (n = 4) and Hubei Beny-like virus (n = 3). Re-annotation of these sequences with BLASTp found evidence of putative homology with hallmark RNA viral genes such as the RNA-dependent RNA polymerase (RdRp; n = 23 sequences), putative major capsid protein (n = 4), viral RNA helicase (n = 4), and polyproteins encoding replicases, including a RNA-dependent RNA polymerase region (n = 5). Moreover, these vHGT sequences had hits to the previously characterized *Symbiodinium* +ssRNA virus (accession KX538960 - KX787934) (Levin et al. 2017).

**Fig. 2:**
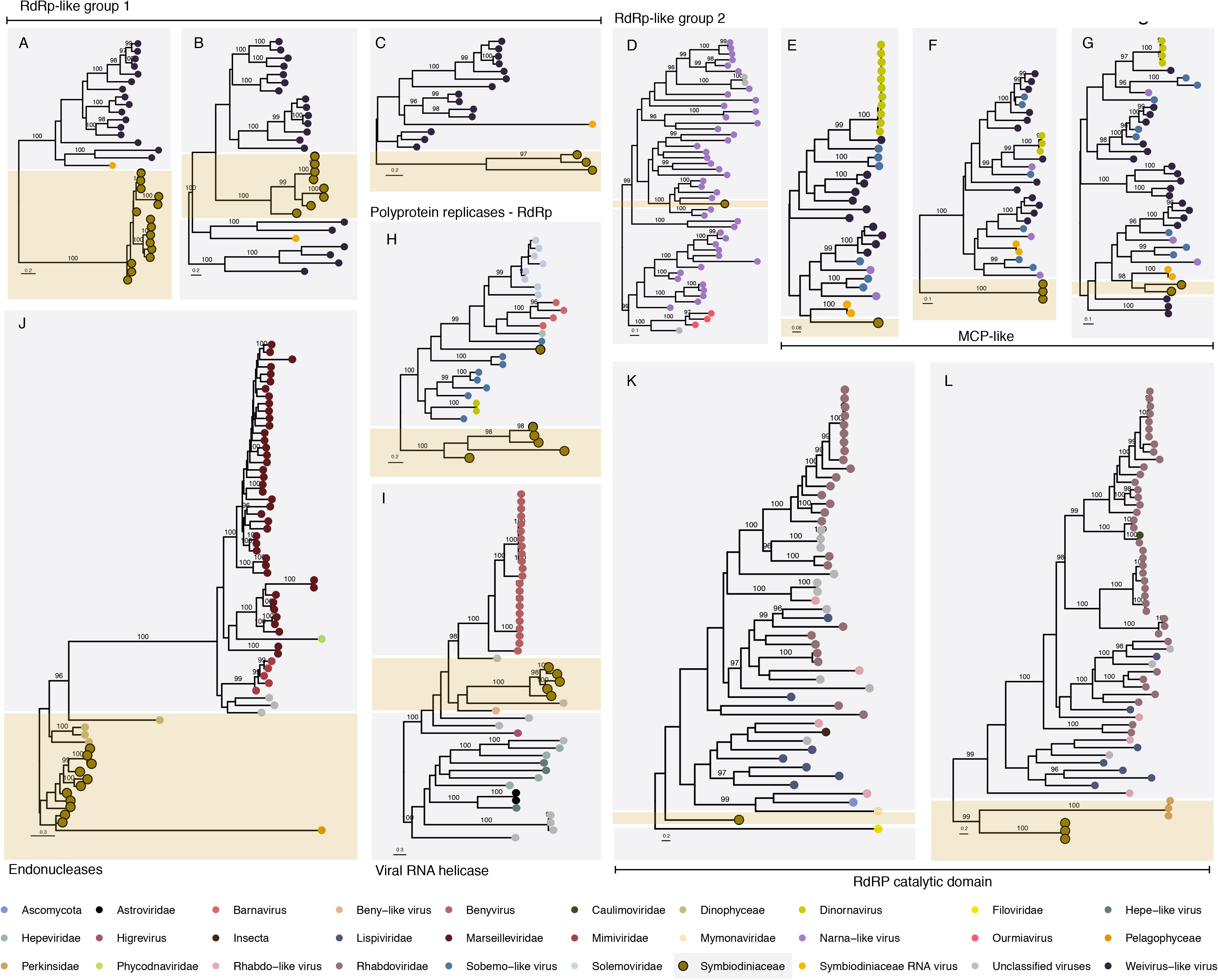
Phylogenetic profile of vHGT’s grouped first by taxonomic affiliation (viral order) and second by putative functional annotation. Unclassified RNA viruses: RNA dependent RNA polymerase (RdRps-like group 1 [A, B, C] and RdRp-like group 2 [D]), Major capsid protein (MCPs-like [E, F, G]), viral RNA helicase (H) and polyprotein coding for replicases including RNA-dependent RNA polymerase region (I); Pimascovirales: restriction endonuclease (Endonuclease) (J); and Mononegavirales: RNA-directed RNA polymerase catalytic domain (RdRP catalytic domain) (K and L). Sequence names are not shown in the trees to enhance readability; the taxonomic affiliation of each sequence in the trees is represented by a colored circle, with the colors described in the legend at the bottom of the figure. Symbiodiniaceae vHGTs are highlighted with thicker borders; only bootstrap support values > 95% (from 10000 bootstrap replicates) are shown. The full phylogenies with sequence names and branch support values are included in Supplementary Fig. S4.

##### 3.4.1.1 RdRp-like phylogenies

The Symbiodiniaceae vHGT-derived RdRps-like sequences were grouped into four phylogenies that matched two different viral groups, indicating that multiple independent vHGTs had occurred. The first viral group (RdRp-like group 1), comprising the sequences from phylogeny A (n = 15 sequences), B (n = 8) and C (n = 3) (Fig. 2 and Supplementary Fig. S4), have hits to Weivirus-like viruses and the *Symbiodinium* +ssRNA virus. In the phylogenetic tree associated with each group, all Symbiodiniaceae vHGTs are clustered into a separate clade or within Weivirus-like viral sequences, distant from the *Symbiodinium* +ssRNA virus. Weivirus-like viruses were first identified in the transcriptomes of molluscs and are related to alveolate EVEs (Shi et al. 2016), and were later found in the Porifera *Halichondria panacea* (breadcrumb sponge) (Waldron et al. 2018).

The second viral group (RdRp-like group 2) comprised the phylogeny D (n = 1), and was a single Symbiodiniaceae sequence from the Bmin genome. This sequence has hits with RdRps from narna-like viruses and is located in a clade containing crustacean and Porifera associated viruses (Beihai and Wenling narna-like viruses and Barns Ness breadcrumb sponge narna−like virus) (Shi et al. 2016). Narnavirus genomes often contain a single open reading frame coding for RdRp. This virus replicates in the cytoplasm of the host and is suggested to be transmitted vertically through cell division, or horizontally during the mating cycle of its fungal host (Dinan et al. 2020). Narnaviruses are associated with trypanosomatids (in which a genomic narnavirus-like element was described (Lye et al.), diatoms (Charon et al. 2021), and red and brown algae (Dinan et al. 2020). Hits were also observed with plant viruses (Ourmiavirus), which have putative origins *via* a chimeric fusion between a fungal narnavirus and other plant viruses (Rastgou et al. 2009).

##### 3.4.1.2 MCP-like phylogenies

The MCP-like vHGT sequences formed three phylogenies: E (n = 1), F (n = 3) and G (n = 2) (Fig. 2 and Supplementary Fig. S4). These sequences matched MCPs from the dinoflagellate HcRNAV Dinornavirus, the invertebrate associated Weivirus−like virus, Sobemo−like virus, narna-like virus, and MCPs from the *Symbiodinium* +ssRNA virus. In the phylogenies E and G, vHGTs are clustered with *Symbiodinium* +ssRNA virus; the sequences from phylogeny F are clustered at the base of the tree. These patterns also suggest multiple independent vHGTs, with high divergence of Symbiodiniaceae MCPs-like sequences, given that viral sequence hits and the tree topology is approximately the same in all three phylogenies.

The remaining vHGTs with unclassified viral hits, from phylogenies H (OG7-OG0009590; n = 6) and I (OG3-OG0006249; n = 5), were re-annotated as viral RNA helicases and “polyprotein coding for replicases including RNA-dependent RNA polymerase region” (respectively). In the phylogeny of the polyprotein-like vHGTs (Phylogeny H), one vHGT sequence (Stri.gene18597) is in a clade with fungal and plant Barnaviruses (Nibert et al. 2018) and Solemoviruses from plant hosts (Sõmera et al. 2021), whereas the other vHGT sequences are at the base of the tree. Interestingly, one sequence in this tree is from the plant *Poinsettia*, which is not only a latent virus, but is suggested to be a hybrid between polerovirus and sobemovirus (aus dem Siepen et al. 2005), which seems to be common in these types of RNA viruses. In the phylogeny I, all Symbiodiniaceae vHGT-derived RNA helicase-like sequences) clustered with a plant virus from the genus *Higrevirus* (Hibiscus green spot virus 2), and with a fungal virus, *Agaricus bisporus* virus 8, forming a bigger clade which includes the aggressive plant disease causing viruses from the genus *Benyvirus* (Gilmer et al. 2017), and Beny-like viruses.

#### 3.4.2 Pimascovirales

The second group of viruses with similarities to vHGTs (n = 11) were all contained within one phylogeny: J (n = 11 sequences) (Fig. 2 and Supplementary Fig. S4). These vHGTs are similar to sequences from the Brazilian marseillevirus (n = 2), Noumeavirus (n = 2), Cannes 8 virus (n = 3), and Kurlavirus BKC-1 (n = 2); all of these taxa are from the Marseilleviridae family of large DNA viruses that infect *Acanthamoeba* protists (Aherfi et al. 2014). Re-annotation of these sequences with BLASTp showed that these sequences are restriction endonucleases, which are suggested to be hotspots for mutations and footprints of mobile elements in other Marseilleviridae (Doutre et al. 2014; Mueller et al. 2017); they also play a role in degradation of host DNA during the early stages of Chlorovirus infection (Agarkova et al. 2006) (Phycodnaviridae; another large DNA virus infecting green algae). Although, in this phylogeny, giant and large viral sequences are overrepresented, such as Marseilleviridae, Mimiviridae, and Phycodnaviridae, we found that vHGTs are clustered at the base of the tree, or with another eukaryote, *Pelagomonas calceolata* (Pelagophyceae). We also find that these sequences occur in the genomes of the dinoflagellate *Polarella glacialis*. This suggests a more complex scenario involving multiple independent and possibly ancient acquisitions, given that almost all genomes from the *Symbiodinium* genus, which is the most ancient in relation to the whole family, contained these sequences.

#### 3.4.3 Mononegavirales

The final group of vHGTs have similarity to Mononegavirales, an order of ssRNA negative-strand viruses. These vHGTs were contained within two phylogenies, K and L (Fig. 2 and Supplementary Fig. S4); four of the vHGTs have top hits to the RNA-directed RNA polymerase catalytic domain. Whereas the best hits of the vHGTs were affiliated with metazoan viruses (Tacheng Tick Virus 6 [n = 3] and Wuchan romanomermis nematode virus 2 [n = 1]), the majority of sequences in the tree were from Rhabdoviridae or Rhabdo-like viruses, which are complex and diverse viruses that include several plant pathogens such as Strawberry crinkle virus and Citrus leprosis virus among others (Dietzgen et al. 2017). Whereas in phylogeny K (OG6-OG0008326) the only vHGT present is outside the main group containing metazoan viruses and the plant soybean leaf-associated negative virus, the phylogeny L (OG1-OG0001147; n = 3), the vHGTs are positioned at the bottom of the tree, clustered with another eukaryote (the perkinsid *Perkinsus olseni*). Because we found hits with the same viral sequences that are split into two phylogenies, it is possible that these were two independent HGT events: one involving the *S. microadriaticum*, Smic, Smic_04-503SCI.03 and Smic_CassKB8 genomes, and the other involving the Snat_CCMP2548 genome.

### 3.5 Sequence composition of Symbiodiniaceae vHGTs sequences

We calculated the coding sequence (CDS) GC-content for vHGTs and for all Symbiodiniaceae CDSs predicted in the genomes that contain vHGTs. That later served as a background for comparative analysis (Fig. 1). Whereas the mean GC-content of the (background) Symbiodiniaceae CDSs ranged from 59.06 % (in *S. microadriaticum* CassKB8) to 51.38 % (in *B. minutum*), the mean GC-content of vHGTs ranged from 60.13 % (in *S. natans* CCMP2548) to 51.29 % (in *B. minutum*). Although the GC-content of the vHGTs was marginally lower than the background CDSs, this difference was statistically significant (*P* = 3.6E-07). However, when these two sets are compared genome-by-genome (i.e., comparing the vHGTs and background CDSs from the same genome) we only found significant GC-content differences in the Smic_CassKB8 (*P =* 1.4E-06), Smic (*P =* 2.4E-05), Snec_CCMP2469 (*P =* 4.4E-05), and Smic_04-503SCI.03 (P= 3.9E-04) genomes. This comparison was not performed for the Cgor, and Slin_CCMP2456 genomes because the number of vHGTs was too small for statistical analysis. There were no significant differences in the length of CDSs between the vHGTs and the background CDSs when comparing all sets together (*P* = 8.0E-02) or individually, although vHGTs are slightly longer (Supplementary Table S3 and S4).

### 3.6 Presence of introns and relic DinoSL motifs in Symbiodiniaceae vHGTs

The vHGTs are predominantly encoded by genes with multiple exons, with only 4/59 present as single-exon genes. The average number of introns per gene in the vHGT gene sets is either roughly equal to (e.g., in *S. microadriaticum* CassKB8), or lower than (e.g., in *S. microadriaticum*) the background gene set for each genome (Fig. 1) (Supplementary Table S6). The average intron length of the vHGT genes tends to differ from that of background genes in each genome, however, this could be a result of the small number of viral HGT genes being analyzed.

Of the 59 vHGT-derived genes, 13 have a DinoSL sequence that is upstream (within 2 kbp) of the first exon (Supplementary Table S7); there was no evidence of relic DinoSL sequences downstream of the identified DinoSL, in close proximity to the vHGT genes.

### 3.7 Distribution of vHGT sequences in Symbiodiniaceae genome scaffolds

We evaluated if there was enrichment of vHGT genes in any of the genome scaffolds in the 10 studied taxa. Our reasoning was that multiple vHGTs in a single scaffold from the same taxonomic source may indicate that they were derived from an exogenous virus that was included during sequencing (i.e., a contaminant in the assembly), or it could represent a complete or near-complete viral genome that had integrated into host DNA. The majority of vHGTs (53 out of 59) were not on the same scaffold as another vHGT. However, in two of the *S. microadriaticum* genomes we found cases of two adjacent vHGTs; these genes were in scaffold3501 from Smic_04-503SCI.03 (gene15955-15956) and in scaffold792 and scaffold67 from Smic (gene19249 -19250 and gene3240-3241, respectively) (Supplementary Table S2). In all three cases, one of the proteins in each pair has similarity to viral RNA-dependent RNA polymerase proteins and the other to viral major capsid proteins, which are the two proteins that comprise the +ssRNA *Symbiodinium* virus genome.

### 3.8 *Symbiodinium* +ssRNA virus similarities in Symbiodininiaceae genomes

We found that the majority of vHGTs (n = 34) had similarity with the most well characterized virus known to infect this group, the *Symbiodinium* +ssRNA virus (TR74740 c13_g1_i1 and _i2, accession KX538960 - KX787934) (Levin et al. 2017; Montalvo-Proaño et al. 2017) that have been implicated in coral holobiont health (Grupstra et al. 2022), and also with the *Heterocapsa circularisquama* RNA virus (HcRNAV; another dinoflagellate infecting-virus). In addition, in two of the *S. microadriaticum* genomes we found adjacent vHGTs re-annotated using BLASTp as the two open reading frames (ORFs) that compose the complete genomes of these two viruses: RNA-dependent RNA polymerase (RdRp) (n = 21), and putative major capsid protein (MCP; n = 4) (Supplementary Table S2). Given these findings, we compared the composition and genomic context of the MCPs-like and RdRp-like vHGTs, with respective sequences from the +ssRNA *Symbiodinium* and HcRNAV viruses. We found that the mean GC-content of Symbiodiniaceae vHGTs MCPs-like and RdRps-like genes was higher (52.95% and 53.76%, respectively) than in the putative +ssRNA *Symbiodinium* virus homologs (48.10% and 43.95%, respectively), lower than the HcRNAV homologs (58.06% and 54.03%, respectively), and lower than in the full Symbiodiniaceae genomes (56.79%). In addition, the mean CDS length of MCPs-like genes was slightly lower than their viral counterparts, whereas the RdRps-like genes were slightly longer (Supplementary Table S4 and S5).

Whereas these proteins do not appear to be expressed (i.e., they did not have any RNA-seq reads aligned using HISAT2; Supplementary Fig. 1A, 2A, and 3A), they were on scaffolds with other expressed multi-exon genes. Realignment of selected RdRp-like and MCP-like proteins against the regions of the host genome that encoded these genes (see Methods) demonstrated that all six of these proteins are significantly degraded (Supplementary Fig. S1-3). These genes often had similarity to only part of the query viral protein (e.g., Supplementary Fig. S1B), in-frame stop codons, or frame-shift mutations (e.g., Supplementary Fig. S2C) that would suggest that they are no longer under selection and have become pseudogenes. Furthermore, two of the putative integrated viral genomes, one from *S. microadriaticum* (gene3240-3241) and one from *S. microadriaticum* 04-503SCI.03 (gene15955-15956) appear to be homologous (Supplementary Fig. S1 and S3), potentially having arisen from an infection in the common ancestor of the two isolates. The third putative integrated genome (gene19249 -19250) only appears in the *S. microadriaticum* genome.

### 3.8 Genomic context of Symbiodiniaceae vHGTs

Through analysis of the position of these sequences in the genome scaffolds, it was observed that the majority of vHGTs encoding MCP-like and RdRp-like genes were located at the start of their scaffolds, or in single gene scaffolds. The CDSs of genes of non-vHGTs up- and downstream of vHGTs (if available) were annotated using the BLASTp online suite; we found that all vHGTs on scaffolds with other genes were located within non-viral host sequences (Supplementary Table S8). In these scaffolds, we found the presence of protein sequences such as the CCHC-type domain-containing protein (n = 2 sequences in total), that was shown in other eukaryotes as either an anti- or pro-viral, targeting several viruses such as RNA viruses (Hajikhezri et al. 2020), that could also be “hijacked” by nuclear-replicating viruses to promote viral production and are also induced after heat shock stress conditions (Younis et al. 2018). Others genes located close to vHGTs are putative E3 ubiquitin-protein ligases (n = 5), RING finger proteins (n = 3; which have roles in viral evasion of host innate immunity) (Xu et al. 2017), F-box proteins (n = 4; which is a component of E3 ubiquitin–ligase complex that targets proteins for ubiquitination and degradation) (R.L. Correa et al. 2013), and the ankyrin repeat domain-containing proteins (n = 6; which are known to regulate virus-host interactions) (Than et al. 2016).

Furthermore, some vHGTs were found to be located in close proximity to transposons, retrotransposons, and repetitive sequences, such as the Retrovirus-related Pol polyproteins (n = 8), Transposon Ty2 Gag-Pol polyprotein (n = 2), pentatricopeptide repeat (n = 2), LINE-1 retrotransposable element ORF2 protein (n = 1), Copia protein (n = 5), and Reverse transcriptase domain-containing protein (n = 2). Finally, three scaffolds were identified in *S. microadriaticum* genomes (Fig. 3) in which the MCPs-like and RdRps-like genes were located adjacent to each other on the same scaffold (scaffold67 and scaffold792 for Smic and scaffold3501 for Smic 04-503SCI.03), where in Smic sequences were in an orientation that was inverted relative to the +ssRNA *Symbiodinium* RNA virus genome (i.e., RdRp upstream of MCP). On scaffold67 these sequences are flanked by a downstream unannotated protein and by a Retrovirus-related Pol polyprotein (from a type-1 retrotransposable element R2). Finally, on scaffold792, there is a reverse transcriptase domain-containing protein at the end of this scaffold.

**Fig. 3:**
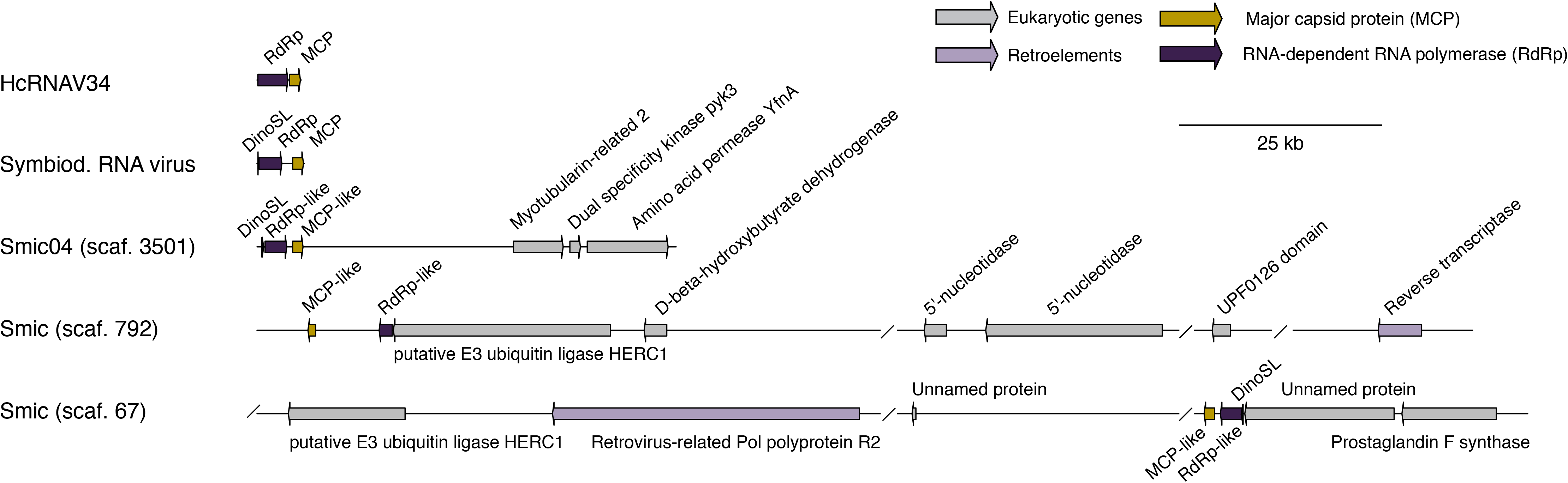
Genome maps comparing the coding sequences of the dinoflagellate virus (dinornavirus) HcRNAV34 (*Heterocapsa* circularisquama RNA virus - accession: AB218608.1) and Symbiod. RNA virus (*Symbiodinium* +ssRNA virus - accession KX538960.1) with the putative integrated viral genomes in *S. microadriaticum* (Smic and Smic04). The putative viral elements (vHGTs) annotated as RdRp-like and MCP-like proteins are shown, as are the genes surrounding these elements. Each feature points in the direction that it is encoded and a scale bar is shown in the top right corner. Slash bars (/) denote intergenic regions. Abbreviations: scaf. (scaffold); DinoSL (dinoflagellate splice leader; annotated as small arrows); RdRp and RdRp-like (RNA dependent RNA polymerase); MCP and MCP-like (major capsid protein); Retrovirus-related Pol polyprotein R2 (Retrovirus-related Pol polyprotein from type-1 retrotransposable element R2); Amino acid permease YfnA (putative amino acid permease YfnA).

## 4 Discussion

In this study, we describe the function, genomic features, and taxonomic distribution of 59 high-confidence viral-derived HGT genes present in 10/13 Symbiodiniaceae genomes. These genes were identified using a combination of HGT scoring metrics, phylogeny-based detection, and significant manual curation. Due to the stringent nature of the filtering that was applied and the lack of a comprehensive database consisting of dinoflagellate-infecting virus genomes, the number of vHGTs we found likely represents a lower bound of the actual number of viral HGT events in these genomes. Nevertheless, by describing a more conservative set of viral HGT candidates in Symbiodiniaceae, we open-up the possibility for additional experimental evaluation to be undertaken to advance functional analysis of these genes. There were between 1-14 vHGTs identified in each genome, comprising ∼0.2% of the total Symbiodiniaceae gene repertoire. Whereas HGT score metrics, such as AI and HGT indexes, are able to identify sequences that are highly similar to foreign sources, phylogenetics can identify more distantly related sequences. Thus, by combining the two approaches we have a higher likelihood of identifying genuine viral HGT events (Fan et al. 2020).

Previous reports of viral genes in Symbiodiniaceae genomes showed that in *S. microadriaticum*, genes with similarity to known viral sequences accounted for < 3% of the total gene inventory (Aranda et al. 2016), and in *C. goreaui* this number was only ∼0.1% (Liu et al. 2018). In the latter case, regions were identified that match viruses known to infect *C. goreaui* (Weynberg et al. 2017) although the origin in these genomes was not clear. Shoguchi et al. (2018) suggested that DNA methylation in some Symbiodiniaceae genomes (clade SymA and SymC) was related to the presence and expansion of endogenous retroviruses such as retrovirus-related genes. In addition, in the *S. microadriaticum* genome, recently expanded Ty1-*copia*-type LTR retrotransposons are activated under acute temperature increase (Chen et al. 2018).

Our findings expand upon these studies by demonstrating the presence of candidate viral sequences from giant virus sources (Pimascovirales) that are potentially functioning as endonucleases, as well as polymerase sequences originating from RNA viruses that could function in viral replication. Exploration of vHGT sequence composition and structural features showed that their GC-contents are slightly lower than for Symbiodiniaceae background CDSs, but higher than homologous viral sequences. The sizes of the coding sequences do not differ from the background Symbiodiniaceae CDS repertoire. This could indicate that some of these transfers were recent in the evolutionary history of this group and are still in the process of being ameliorated into the recipient host genome (Lawrence and Ochman 1997). Moreover, we found that viral sequences are primarily localized to different scaffolds in the genome but tend to be near eukaryotic genes with known roles in virus-host interaction. Finally, we found these viral sequences near mobile genetic elements, such as retrotransposons and transposons, that may act not only as preferential integration sites, but as a mechanism for their integration and expansion.

Currently, the best characterized virus that infects Symbiodiniaceae is a positive single-stranded RNA virus (+ssRNA) (NCBI:txid1909300) (Nagasaki et al. 2005; A.M.S. Correa et al. 2013; Montalvo-Proaño et al. 2017; Weynberg et al. 2017), first described in stressed *Montastraea cavernosa* coral viromes (A.M.S. Correa et al. 2013), that is related to the dinoflagellate virus *Heterocapsa circularisquama* (HcRNAV) from the *Alvernaviridae* family. Its complete genome was described by Levin et al (2016); they found that thermally stressed *Cladocopium* (formerly *Symbiodinium* clade C1) populations experienced an extreme +ssRNA virus infection, suggesting that viral infections may influence *Symbiodinium* thermal sensitivity and consequently, coral bleaching. Nevertheless, these authors were unable to amplify MCP sequences from genomic DNA (only from cDNA), suggesting that these sequences were not endogenous in the host genome, but rather exogenous. Moreover, they reported the presence of a DinoSL at the 5’-end of the amplified viral transcripts, proposing that this motif was incorporated into the viral genome as a form of “molecular mimicry” to evade the host immune response and to drive efficient translation of the viral genome by the host cellular machinery.

It has been suggested that viral infections in Symbiodiniaceae are latent, given that VLPs and expressed sequences were found in asymptomatic and healthy cultures subjected to stress conditions (Wilson et al. 2001; Lawrence et al. 2014; Brüwer et al. 2017). Lawrence et al. (2017) showed that in Symbiodiniaceae cultures exposed to UV stress, there was upregulation of virus-like genes associated with viral genome replication and protein production, such as ubiquitin-encoding genes and homing endonucleases that could be involved in the excision of the integrated viral genome. This pattern is consistent with latent virus replication and agrees with our results.

Latent viruses are well described in plants and often do not have any apparent symptoms in wild populations, however, they can cause disease in crops or due to experimentally created stresses such as mechanical wounding (Roossinck and García-Arenal 2015). Mixed viral infections and changes in environmental conditions trigger the switch from latent to active infections and diseases (Takahashi et al. 2019). In plants, the integration of a virus genome is not required for latent viral replication, although partial genome integration and endogenization of entire viral genomes may play a major role in latency (Takahashi et al. 2019). Latent infections and EVEs present major risks for cultivated crops if they become active (Bertsch et al. 2009). This is the case for the hybrid *Musa* (Iskra-Caruana et al. 2010), where the endogenous *banana streak virus* (eBSV), which has multiple copies integrated into the host genome, becomes activated under stress, recombining, producing episomes, viral particles, and disease symptoms such as streaks and necrosis in plant leaves. Recent work using CRISPR/Cas9-mediated editing of eBSV has been successful in preventing activation of these viruses (Tripathi et al. 2019). In contrast, benefits of latent viruses and EVEs are heat and drought tolerance, protection of host plants against virulent viral strains (Agüero et al. 2018), and suppression of heterologous viral infections (Staginnus et al. 2007).

### 4.1 Proposed models for vHGTs in Symbiodiniaceae genomes

Given our findings, we propose a model with two hypothetical scenarios to explain the occurrence of viral sequences in the genomes of Symbiodiniaceae and the underlying mechanisms for their integration (Fig. 4). In this model, whereas several viral groups infect Symbiodiniaceae, independently entering the host cell and subjugating the host machinery to produce viral mRNAs, some virus sequences become endogenous viral elements (EVEs) by a series of incidental integration events into Symbiodiniaceae genomes (A). For this to occur, viral mRNAs need to be reverse transcribed before they can be integrated into the host genome. This process might occur *via* the mRNA recycling mechanism that is prevalent in dinoflagellates (Stephens et al. 2020), whereby spliced-leader-containing transcripts are reverse transcribed (possibly mediated by retrotransposons (Lee et al. 2014; Song et al. 2017) into DNA before being integrated back into the genome through non-homologous recombination (Slamovits and Keeling 2008). The infecting viral elements must escape degradation by host defenses. Because viral mRNAs are highly expressed during an active infection, the host spliceosomal machinery may erroneously add a DinoSLs motif onto viral transcripts, making the transcripts appear “native”. This may have allowed the viral genome to better evade host defenses and providing the virus with a dinoflagellate TATA-box-like promoter sequence; dinoflagellates appear to use a TTTT motif (which is also present in the DinoSL) in place of the canonical TATA motif in other eukaryotes (Guillebault et al. 2002). This is supported by the presence of a DinoSL in the *Symbiodinium* RNA virus genome (Levin et al. 2017) and suggests that this motif could have been acquired by the virus from the host either in the common ancestor of these viruses or during each new infection cycle. Whereas convergence is also possible (i.e., independent evolution of a DinoSL-like motif), it is less likely given the high level of sequence similarity between the viral and host DinoSL sequences. Furthermore, the occurrence of DinoSL sequences upstream of 13 vHGTs, plus the localization of these sequences near retrovirus related polyproteins, copia proteins, and proteins containing reverse transcriptase domains in the Symbiodiniaceae scaffolds, indirectly support this mechanism of integration.

**Fig. 4:**
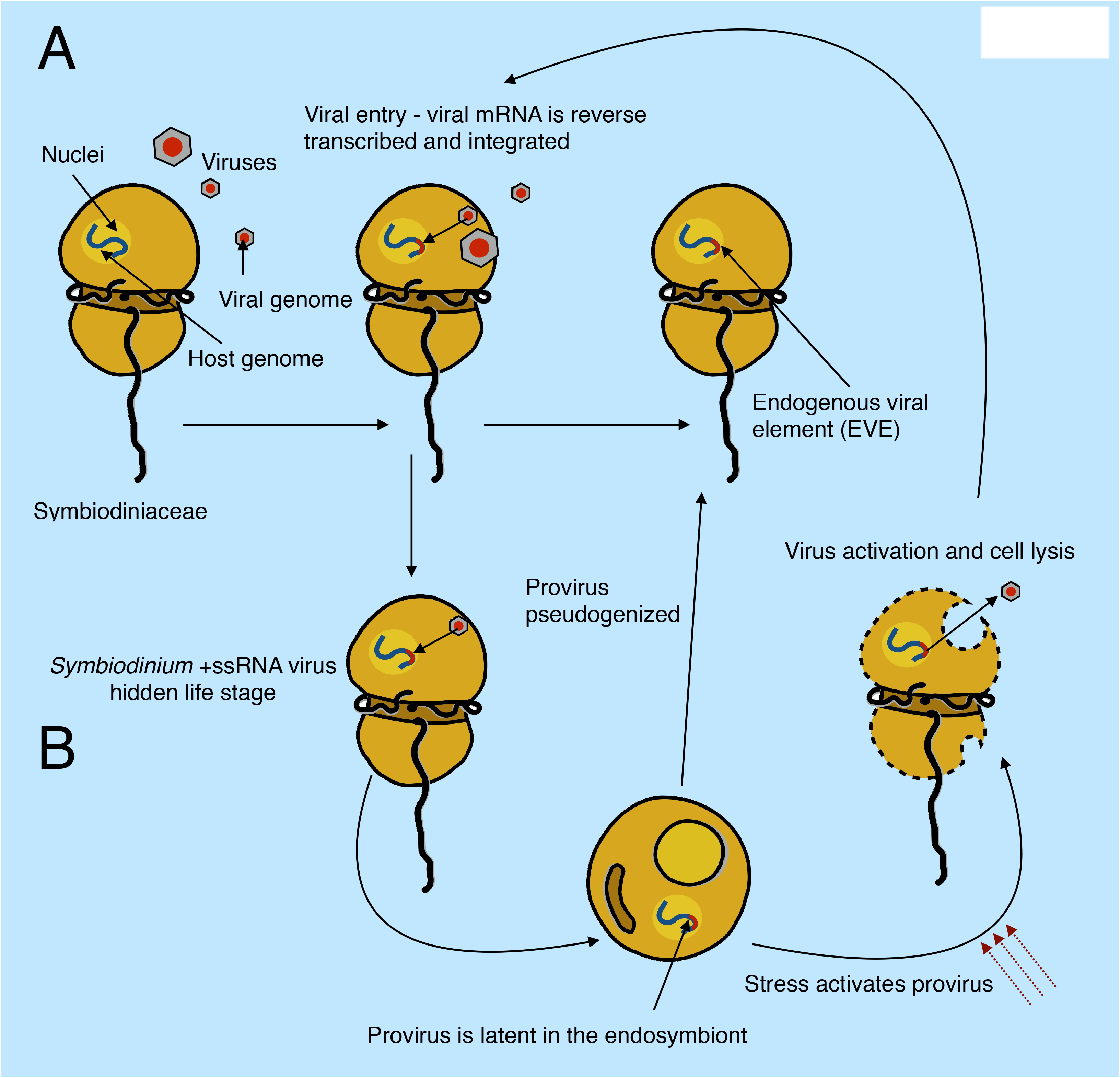
hypothetical model with two scenarios (A and B) explaining the origin of vHGTs in Symbiodiniaceae genomes. During an acute infection, viral particles enter the host cell and viral mRNAs accumulate in the cytoplasm to a high abundance. Host machinery can accidentally incorporate viral mRNA into its genome *via* reverse transcription and non-homologous recombination; this may use the same (or similar) mechanisms as the mRNA recycling process that is prevalent in dinoflagellates. These viral sequences, known as endogenous viral elements (EVEs), would be “dead on arrival” and are expected to decay into pseudogenes (which we would then observe in the genome as vHGTs). In the second scenario (B), integration of the virus genome into the host genome is part of a previously unobserved transient life stage of these viruses (specifically the *Symbiodinium* +ssRNA virus). The virus infects and is incorporated into the host genome *via* the same mechanisms as the first scenario (A) however, instead of this process resulting in non-functional EVEs it produces active proviral elements. These integrated viral genomes can remain silent during times of low host abundance (e.g., during the free-living stage of facultative Symbiodiniaceae lifecycles) and can become activated by environmental cues (such as stress or high Symbiodiniaceae cell density) that result in viral lysis, escape from the host cell and new infections in susceptible hosts. The proviral elements can also become deactivated, either through random mutation or host driven process, resulting in EVEs that are expected to decay into pseudogenes, which we would then observe in the genome as vHGTs.

However, given that we found complete viral genomes with similarity to the +ssRNA *Symbiodinium* virus, we propose a second scenario (B) whereby vHGTs from +ssRNA *Symbiodinium* viruses resulted from a previously hidden (cryptic) life stage of these viruses. In this scenario, copies of viral genomes become endogenized and deactivated by the host genome, decaying as pseudogenes. Alternatively, they remain intact, transcriptionally silent (as proviruses), and may eventually become activated in response to stress, shifting from a proviral silent life-stage to being infectious. These RNA viruses could actively exploit the dinoflagellate mRNA recycling process to facilitate their integration into host DNA although this would require the virus to also have a system for permanence (survival) and activation, both of which are unknown at this time. However, given that the genes associated with the three putative integrated *Symbiodinium* +ssRNA virus genomes, identified in two of the *S. microadriaticum* isolates, appear to have become pseudogenized, it seems likely that many of the viral genes that we identified in the Symbiodiniaceae genomes are (or are becoming) pseudogenes. This is an expected result given that there is strong selective pressure on the host to inactivate endogenous viruses and that we only expect to see remnants of inactivated endogenous viruses originating from past (failed) infections in the genomes of extant lineages (i.e., active endogenous viral elements would likely result in an active viral infection that would cause the lineage to perish). In addition, the two vHGT genes that are part of the putative *Symbiodinium* +ssRNA viruses identified in the *S. microadriaticum* (Smic) genome are encoded in the reverse order to what is observed in the available virus reference genome. This may result from the host genome inactivating the invasive viruses by shuffling their gene order, preventing the formation of infective viral particles. Furthermore, the putative inactivation of these elements by the host suggests that they were active when they were integrated into the genome, lending support to the second scenario (which produces viral elements that are “dead on arrival”). Moreover, the multiple, complete and incomplete copies of these viral genomes in Symbiodiniaceae, from different isolates and genera, suggests that viral endogenization is an ongoing process in these organisms, with a potentially rapid turnover of invasion and pseudogenization or activation and infection, similar to Mavirus virophage infection-integration cycles (Hackl et al. 2021).

Under this scenario, endogenization does not need to be an obligatory step in the life cycle of the virus (i.e., it is a transient stage) and does not need to become fixed in all hosts occurring, for instance, only when there is a low number of susceptible hosts. Becoming endogenized when the density of host cells is low is advantageous for the virus, given that a copy of the virus genome is always transmitted to host progeny, effectively preserving the virus until conditions promote the lytic life stage. The virus life cycle may therefore mirror that of the host, remaining inactive during phases where the density of susceptible cells is low, such as during the host coccoid endosymbiont life stage, and becoming active when the density of susceptible cells is high, such as during the host mastigote, free-living stage. From this perspective, the more susceptible stage would be the free-living state with no endogenous provirus in its genome. From the perspective of the Symbiodiniaceae host, it could be argued that having an endogenous provirus would confer superinfection immunity against other exogenous viruses *via* cross protection and may benefit the cell under certain conditions (Mette et al. 2002). A hidden transient viral life cycle, if confirmed for these viruses, would unify the current knowledge of viruses in Symbiodiniaceae, explaining the observations of: (1) latent viral infections in healthy algal cultures after stress induction, (2) exogenous RNA viruses that infect Symbiodiniaceae, and (3) our findings of endogenous versions of these exogenous *Symbiodinium* RNA virus genomes.

As previously noted, the triggers that cause the switch from latent to active viral infections and diseases in plants and algae (including cultured Symbiodiniaceae) overlaps with those that trigger and influence coral bleaching and other coral diseases (Harvell et al. 1999; Lesser and Farrell 2004; Randall and van Woesik 2015; Lawrence et al. 2017). It has been recently proposed that breakdown of the Symbiodiniaceae-coral symbiosis due to heat stress (coral bleaching) is caused by increased proliferation of the symbiont (Rädecker et al. 2021). The observed increase in viral particle production under thermal stress may coincide with the increased proliferation of the symbionts during the early stages of coral bleaching. This result suggests that the virus life cycle could be directly linked to that of the host.

Consequently, the main implication of our work is that if EVEs and latent proviruses that are hidden in Symbiodiniaceae genomes could become activated by environmental stressors, that they may cause local outbreaks and possibly contribute to the onset of some coral diseases. To conclude, by describing a variety of acquired viral genes originating from multiple HGT events, we provide strong evidence that viruses have invaded and introduced novel genetic material into Symbiodiniaceae genomes. Our work suggests the possibility that endogenous and latent viruses could modulate and participate in the life cycle, evolution, and potentially breakdown of the Symbiodiniaceae-host symbiosis.

## Data availability

All Symbiodiniaceae sequence data used in this study are available on https://espace.library.uq.edu.au/view/UQ:f1b3a11 and https://espace.library.uq.edu.au/view/UQ:8279c9a. Code to generate the HGT calculations is available at https://github.com/LFelipe-B/Symbiodiniaceae_vHGT_scripts.

## Supplementary data

Supplementary data is available at Virus Evolution online.

## Acknowledgements

LFB wish to thank Paulo Sérgio Salomon (Universidade Federal do Rio de Janeiro - UFRJ) and coordinators from “Programa de Pós-graduação em Biodiversidade e Biologia Evolutiva” (PPGBBE - UFRJ), for the initial conceptual support in early stage of these ideas; Lílian Caesar for critical reading, discussions, love and support during all stages of this work; François Bucchini for support in initial codes; GenoToul bioinformatics platform for access to computing cluster facilities during early stages of this work; Girish Beedessee and Cheong Xin Chan for guidance in data retrieval.

## Conflict of interest

The authors declare no conflict of interest.

Supplementary Fig. S1: Putative integration of an +ssRNA *Symbiodinium* virus genome into the genome of *S. microadriaticum* 04-503SCI.03 (scaffold: Smic_04-503SCI.03.scaffold3501; genes: gene15955 and gene15956). (A) Image taken from IGV showing the region of the genome encoding the two vHGT genes (represented as blue bars; solid bars represent exons and lines represent introns) along with aligned RNA-seq data (gray discontinuous bars). (B) Alignment (generated using exonerate) of a RNA-dependent RNA polymerase protein (YP_009337004.1) against a region (Smic_04-503SCI.03.scaffold3501:13547-15709) of the *S. microadriaticum* 04-503SCI.03 genome corresponding to the vHGT gene15955. (C) Alignment (generated using exonerate) of a major capsid protein (AOG17586.1) against a region (Smic_04-503SCI.03.scaffold3501:16298-17334) of the *S. microadriaticum* 04-503SCI.03 genome corresponding to the vHGT gene15956. In B and C, a predicted intron is highlighted using a blue box and in-frame stop codons using red boxes.

Supplementary Fig. S2: Putative integration of an +ssRNA *Symbiodinium* virus genome into the genome of *S. microadriaticum* (scaffold: scaffold792; genes: gene19249 and gene19250). (A) Image taken from IGV showing the region of the genome encoding the two vHGT genes (represented as blue bars; solid bars represent exons and lines represent introns) along with aligned RNA-seq data (gray discontinuous bars). (B) Alignment (generated using exonerate) of a major capsid protein (AOG17586.1) against a region (Smic.scaffold792:89753-90508) of the *S. microadriaticum* genome corresponding to the vHGT gene19249. (C) Alignment (generated using exonerate) of a RNA-dependent RNA polymerase protein (YP_009342067.1) against a region (Smic.scaffold792:96851-98156) of the *S. microadriaticum* genome corresponding to the vHGT gene19250. In B and C, a predicted intron is highlighted using a blue box, in-frame stop codons using red boxes, and frame-shifts using orange boxes.

Supplementary Fig. S3: Putative integration of an +ssRNA *Symbiodinium* virus genome into the genome of *S. microadriaticum* (scaffold: scaffold67; genes: gene3240 and gene3241). (A) Image taken from IGV showing the region of the genome encoding the two vHGT genes (represented as blue bars; solid bars represent exons and lines represent introns) along with aligned RNA-seq data (gray discontinuous bars). (B) Alignment (generated using exonerate) of a major capsid protein (AOG17586.1) against a region (Smic.scaffold67:1208081-1209117) of the *S. microadriaticum* genome corresponding to the vHGT gene3240. (C) Alignment (generated using exonerate) of a RNA-dependent RNA polymerase protein (YP_009337004.1) against a region (Smic.scaffold67:1209706-1211868) of the *S. microadriaticum* genome corresponding to the vHGT gene3241. In B and C, a predicted intron is highlighted using a blue box, in-frame stop codons using red boxes, and frame-shifts using orange boxes.

Supplementary Fig. S4: Phylogenies organized by grouping (A-L), viral order (Unclassified RNA viruses, Pimascovirales and Mononegavirales) of subject hits and predicted annotation, showing complete sequence names and annotation with branch labels and associated ultrafast bootstrap values.

