## Supplementary Fig. S4 for "Multiple waves of viral invasions in Symbiodiniaceae algal genomes"

#### Unclassified RNA viruses

##### RdRp-like group 1 (Phylogeny A)

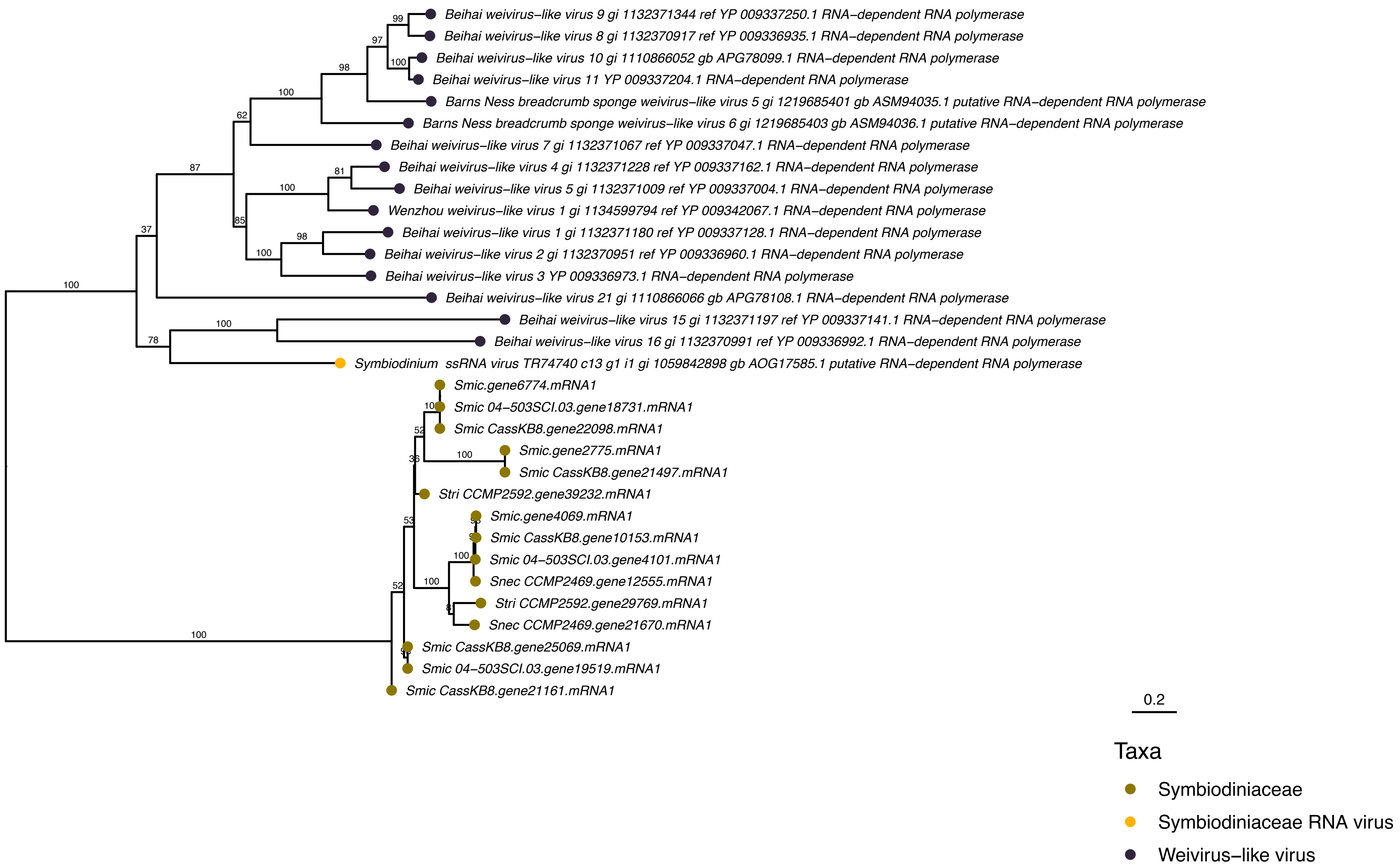


##### RdRp-like group 1 (Phylogeny B)

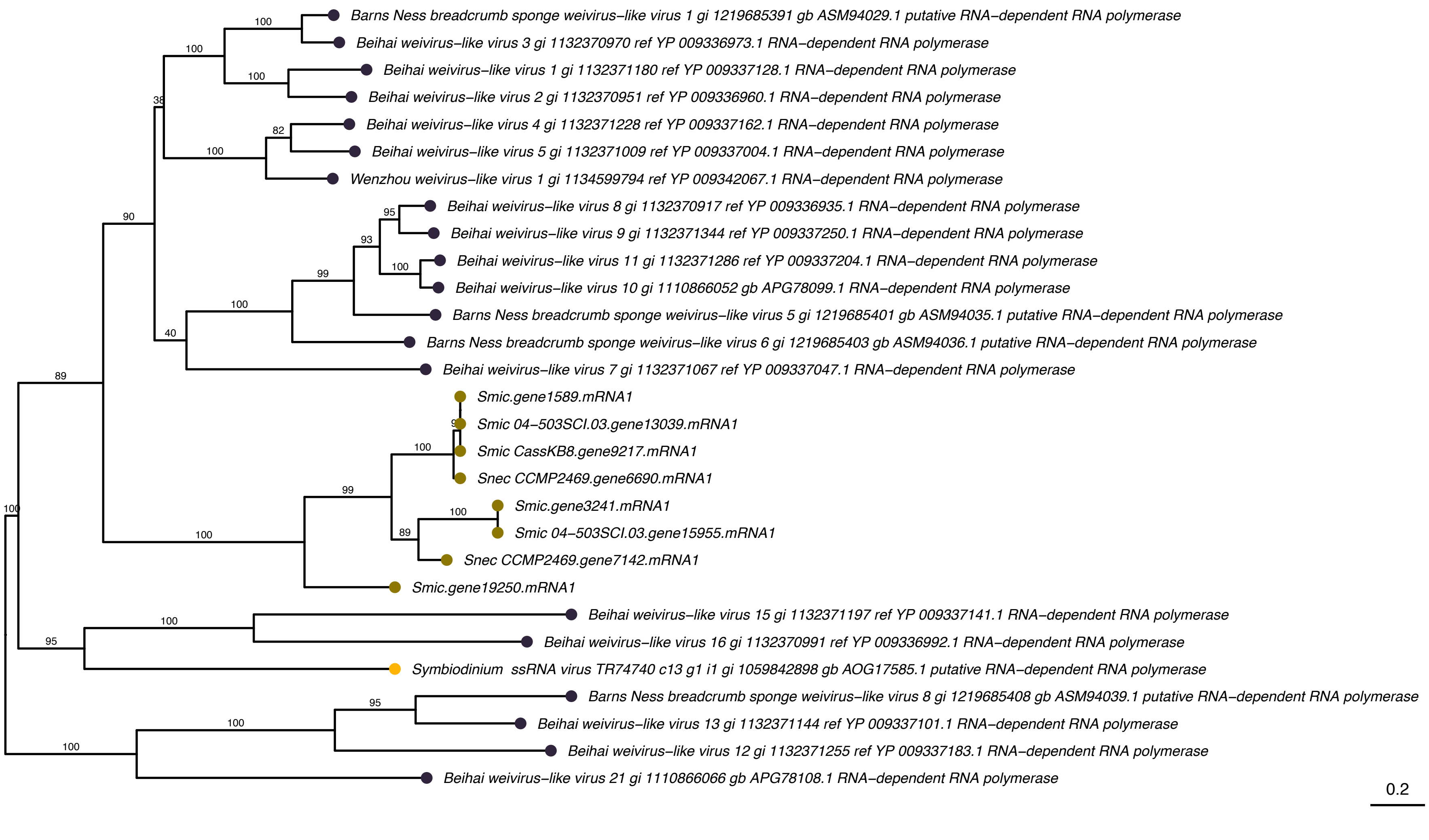

#### Unclassified RNA viruses

#### RdRp-like group 1 (Phylogeny C)

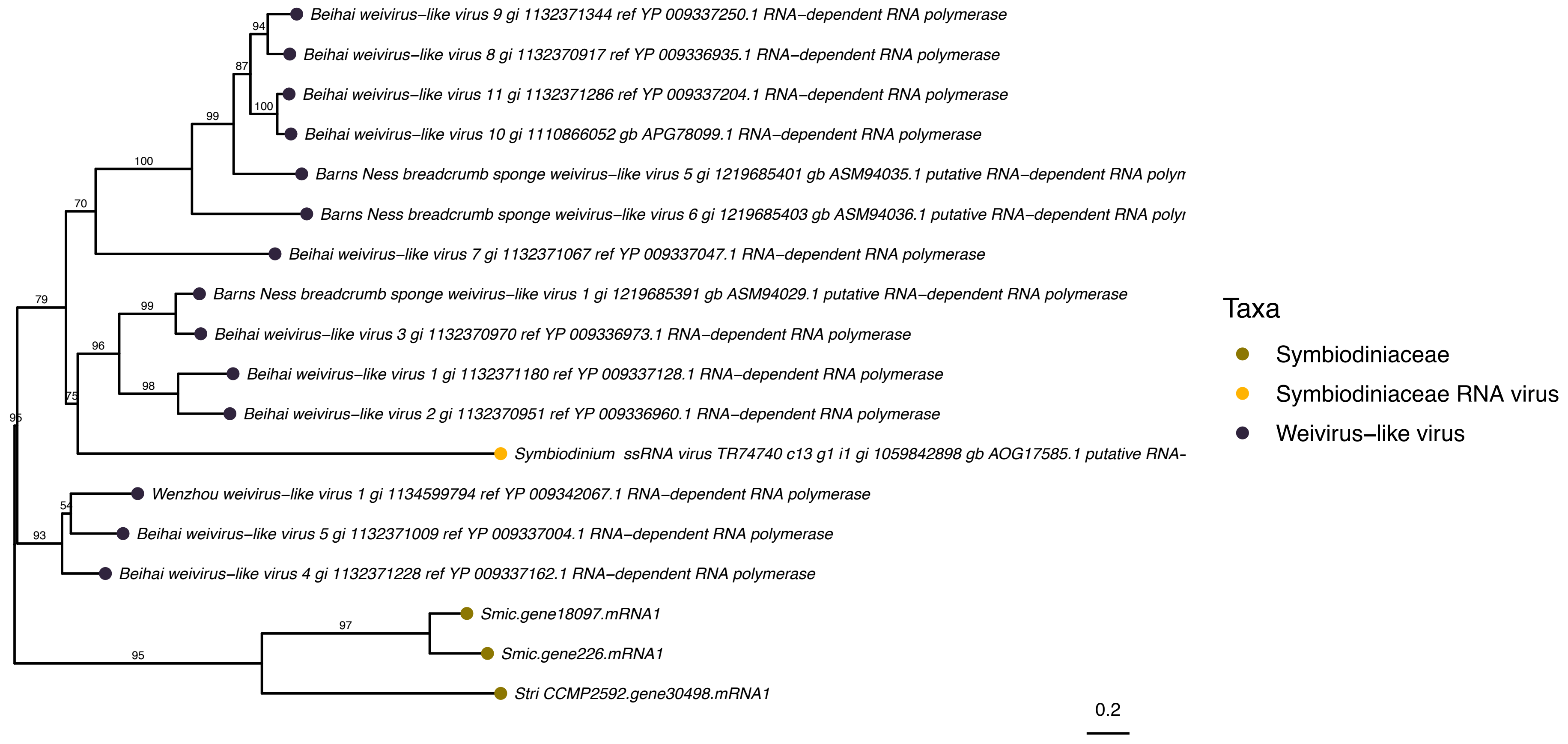


#### RdRp-like group 2 (Phylogeny D)

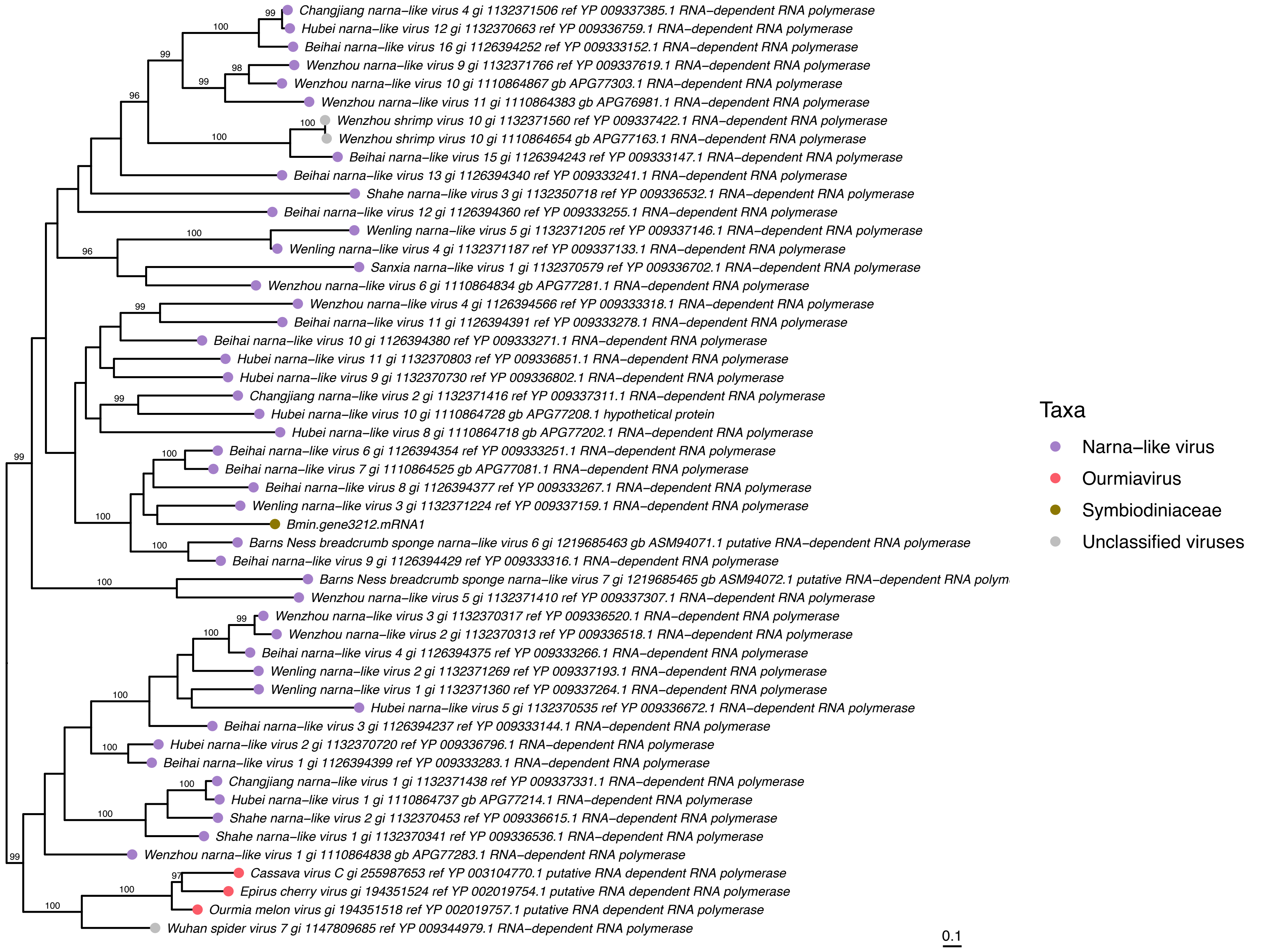

### Unclassified RNA viruses

#### MCP-like (Phylogeny E)

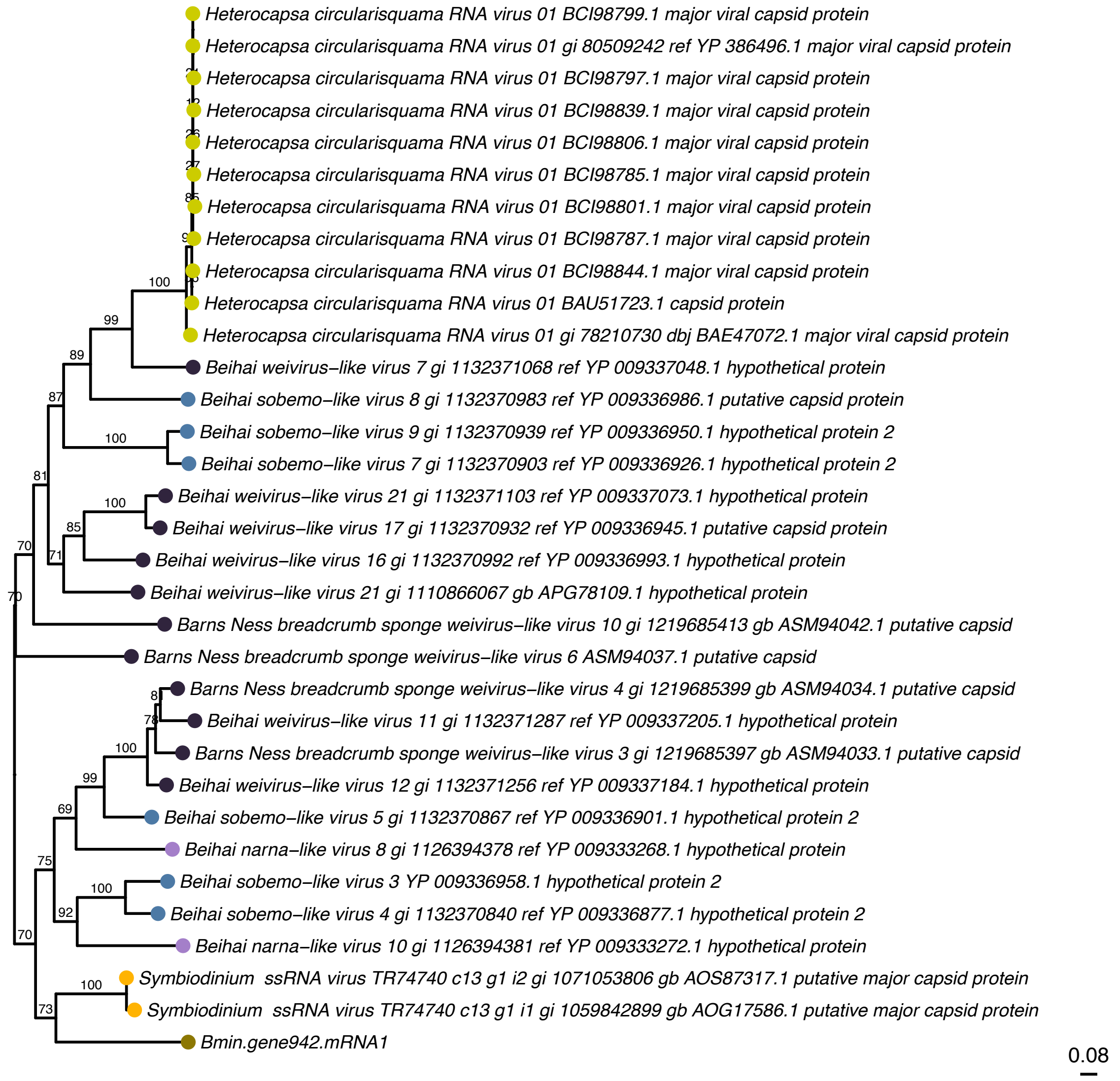


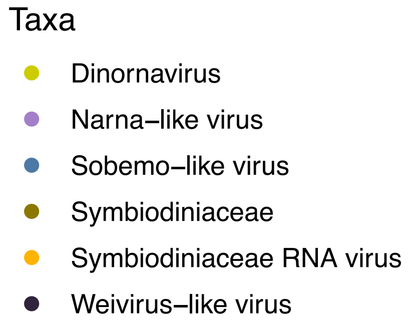


#### MCP-like (Phylogeny F)

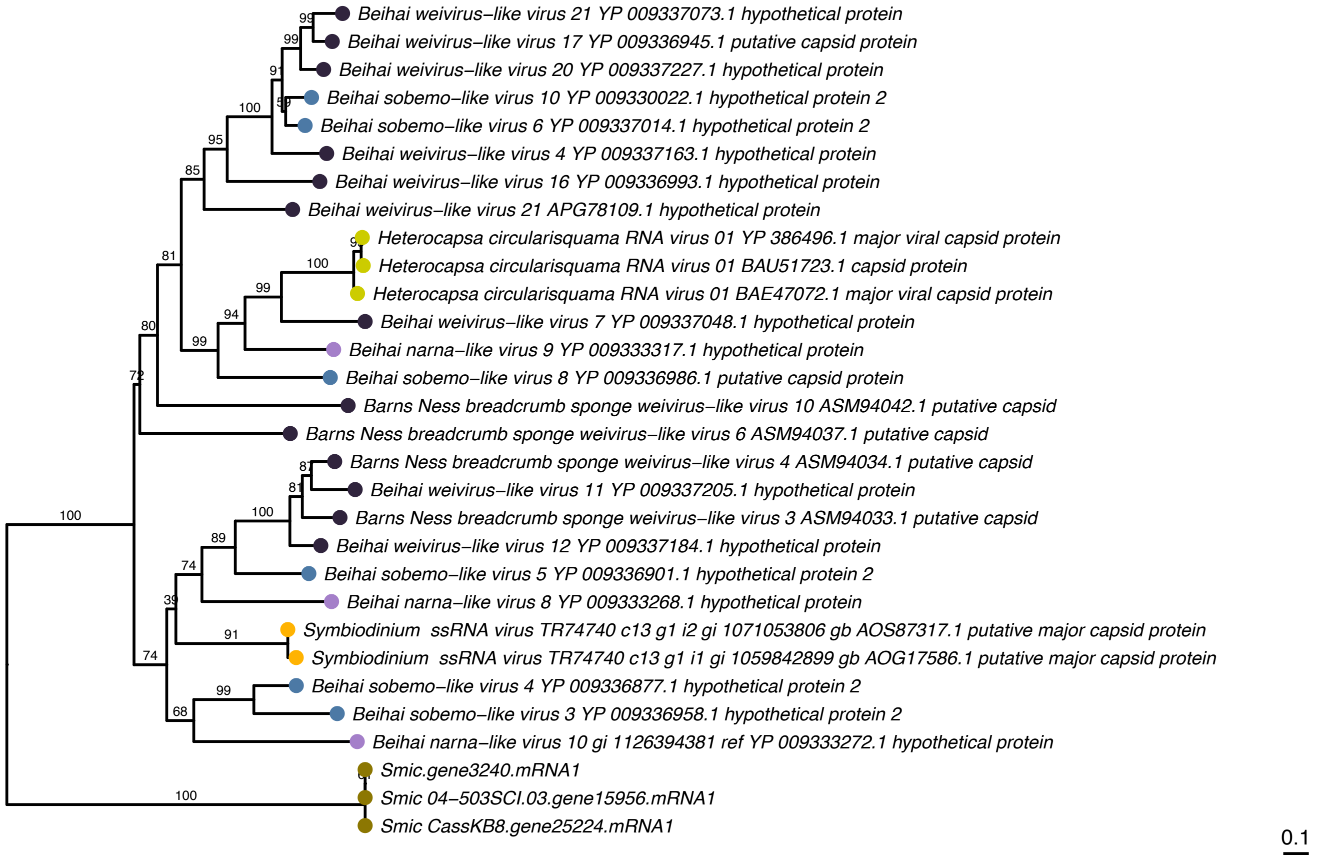

Unclassified RNA viruses

#### MCP-like (Phylogeny G)

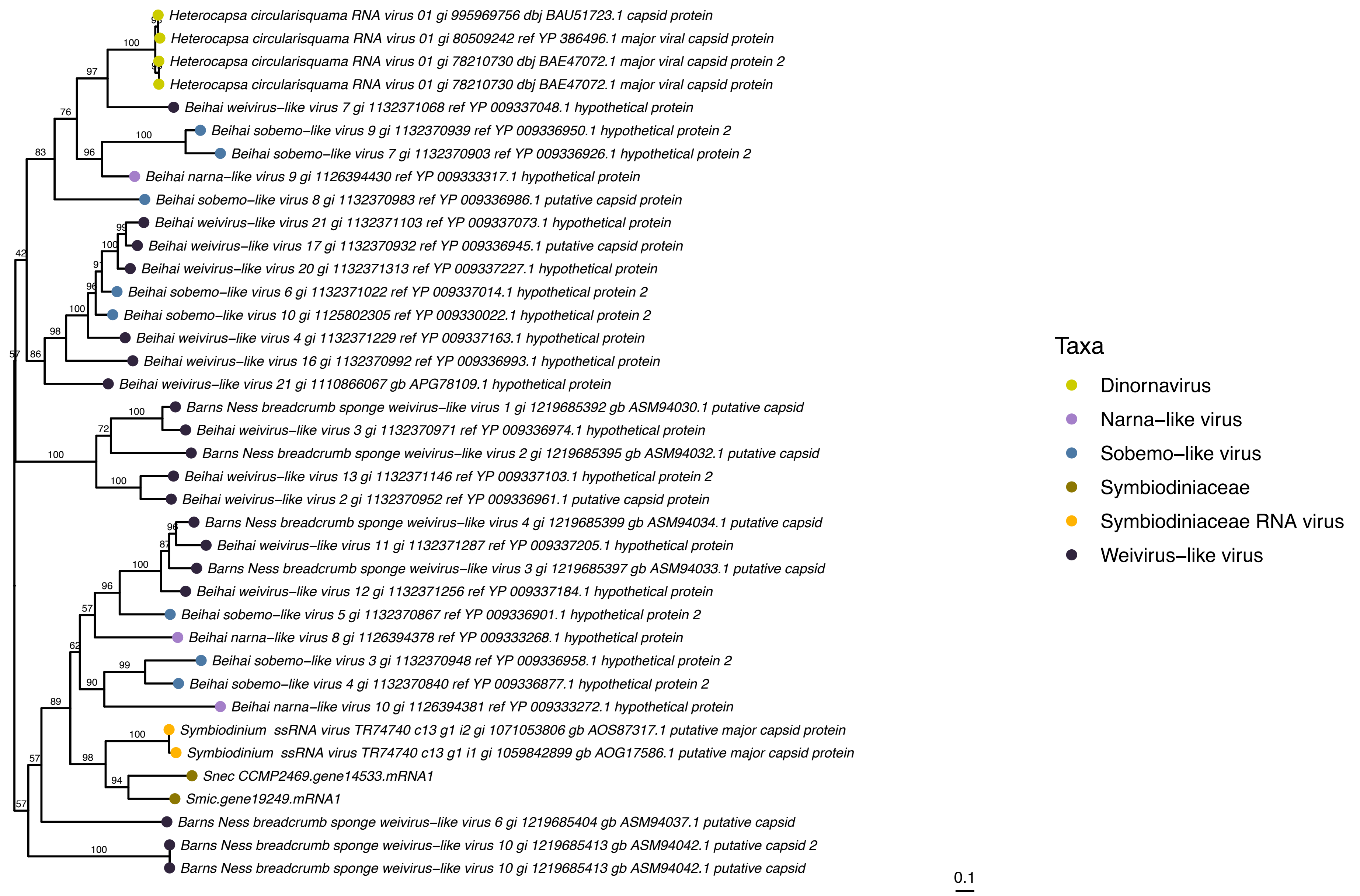


#### Polyprotein replicases - RdRp (Phylogeny H)

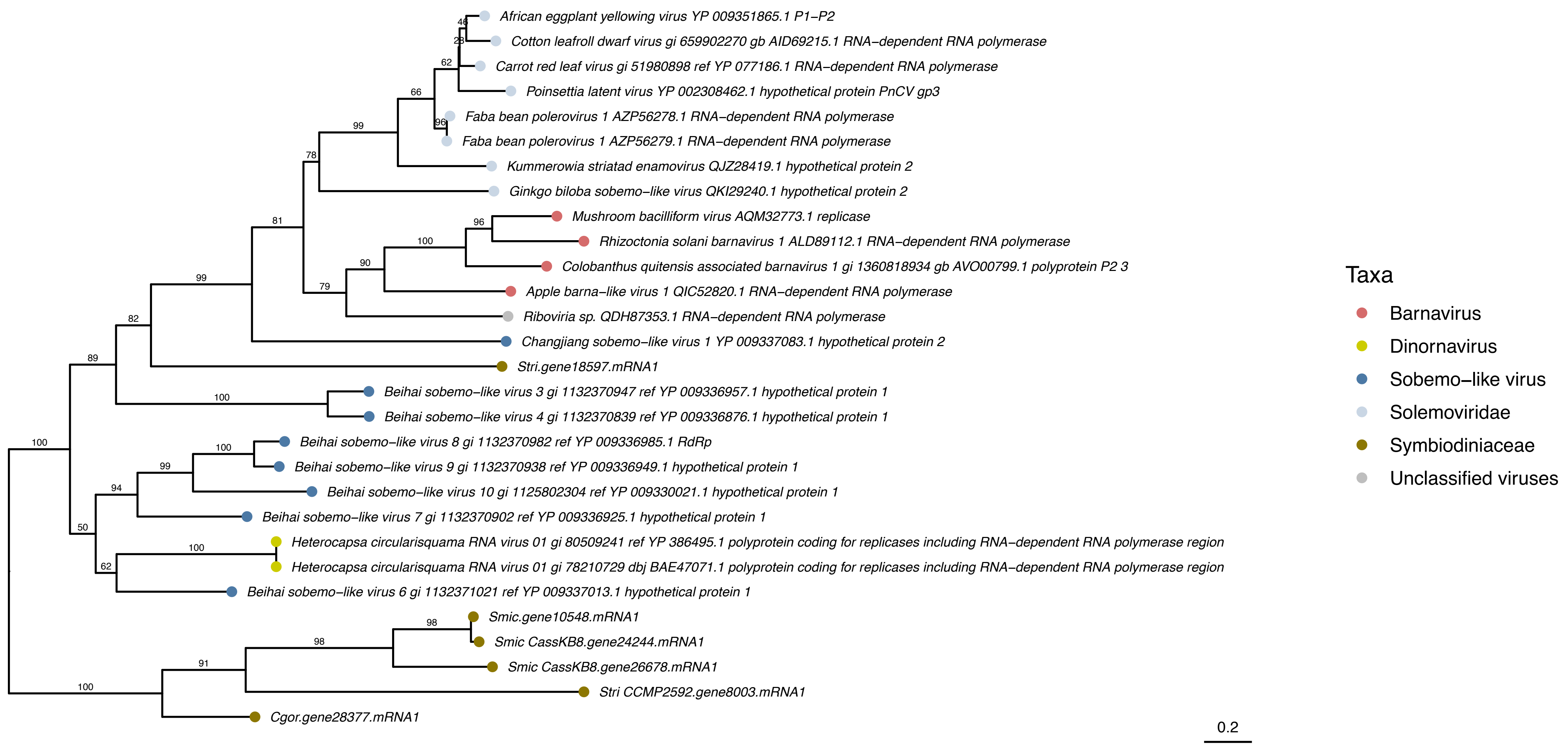

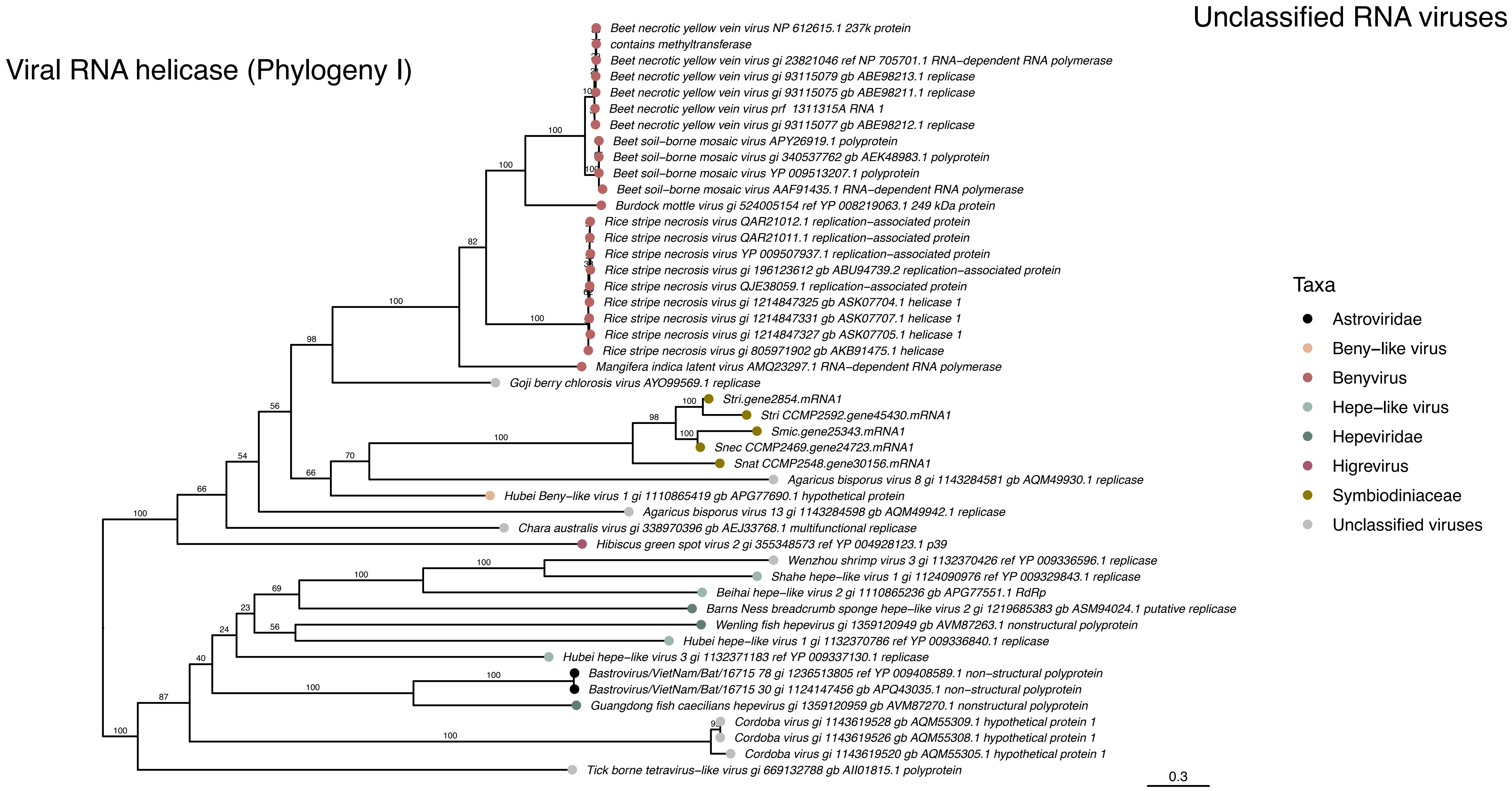
Unclassified RNA viruses
Viral RNA helicase (Phylogeny I)
Beet necrotic yellow vein virus NP 612615.1 237k protein
contains methyltransferase
Beet necrotic yellow vein virus gi 23821046 ref NP 705701.1 RNA–dependent RNA polymerase
Beet necrotic yellow vein virus gi 93115079 gb ABE98213.1 replicase
Beet necrotic yellow vein virus gi 93115075 gb ABE98211.1 replicase
Beet necrotic yellow vein virus prf 1311315A RNA 1
Beet necrotic yellow vein virus gi 93115077 gb ABE98212.1 replicase
Beet soil–borne mosaic virus APY26919.1 polyprotein
Beet soil–borne mosaic virus gi 340537762 gb AEK48983.1 polyprotein
Beet soil–borne mosaic virus YP 009513207.1 polyprotein
Beet soil–borne mosaic virus AAF91435.1 RNA–dependent RNA polymerase
Burdock mottle virus gi 524005154 ref YP 008219063.1 249 kDa protein
Rice stripe necrosis virus QAR21012.1 replication–associated protein
Rice stripe necrosis virus QAR21011.1 replication–associated protein
Rice stripe necrosis virus YP 009507937.1 replication–associated protein
Rice stripe necrosis virus gi 196123612 gb ABU94739.2 replication–associated protein
Rice stripe necrosis virus QJE38059.1 replication–associated protein
Rice stripe necrosis virus gi 1214847325 gb ASK07704.1 helicase 1
Rice stripe necrosis virus gi 1214847331 gb ASK07707.1 helicase 1
Rice stripe necrosis virus gi 1214847327 gb ASK07705.1 helicase 1
Rice stripe necrosis virus gi 805971902 gb AKB91475.1 helicase
Mangifera indica latent virus AMQ23297.1 RNA–dependent RNA polymerase
Goji berry chlorosis virus AYO99569.1 replicase
Stri.gene2854.mRNA1
Stri CCMP2592.gene45430.mRNA1
Smic.gene25343.mRNA1
Snec CCMP2469.gene24723.mRNA1
Snat CCMP2548.gene30156.mRNA1
Agaricus bisporus virus 8 gi 1143284581 gb AQM49930.1 replicase
Hubei Beny–like virus 1 gi 1110865419 gb APG77690.1 hypothetical protein
Agaricus bisporus virus 13 gi 1143284598 gb AQM49942.1 replicase
Chara australis virus gi 338970396 gb AEJ33768.1 multifunctional replicase
Hibiscus green spot virus 2 gi 355348573 ref YP 004928123.1 p39
Wenzhou shrimp virus 3 gi 1132370426 ref YP 009336596.1 replicase
Shahe hepe–like virus 1 gi 1124090976 ref YP 009329843.1 replicase
Beihai hepe–like virus 2 gi 1110865236 gb APG77551.1 RdRp
Barns Ness breadcrumb sponge hepe–like virus 2 gi 1219685383 gb ASM94024.1 putative replicase
Wenling fish hepevirus gi 1359120949 gb AVM87263.1 nonstructural polyprotein
Hubei hepe–like virus 1 gi 1132370786 ref YP 009336840.1 replicase
Hubei hepe–like virus 3 gi 1132371183 ref YP 009337130.1 replicase
Bastrovirus/VietNam/Bat/16715 78 gi 1236513805 ref YP 009408589.1 non–structural polyprotein
Bastrovirus/VietNam/Bat/16715 30 gi 1124147456 gb APQ43035.1 non–structural polyprotein
Guangdong fish caecilians hepevirus gi 1359120959 gb AVM87270.1 nonstructural polyprotein
Cordoba virus gi 1143619528 gb AQM55309.1 hypothetical protein 1
Cordoba virus gi 1143619526 gb AQM55308.1 hypothetical protein 1
Cordoba virus gi 1143619520 gb AQM55305.1 hypothetical protein 1
Tick borne tetravirus–like virus gi 669132788 gb AII01815.1 polyprotein
0.3
Taxa
Astroviridae
Beny–like virus
Benyvirus
Hepe–like virus
Hepeviridae
Higrevirus
Symbiodiniaceae
Unclassified viruses

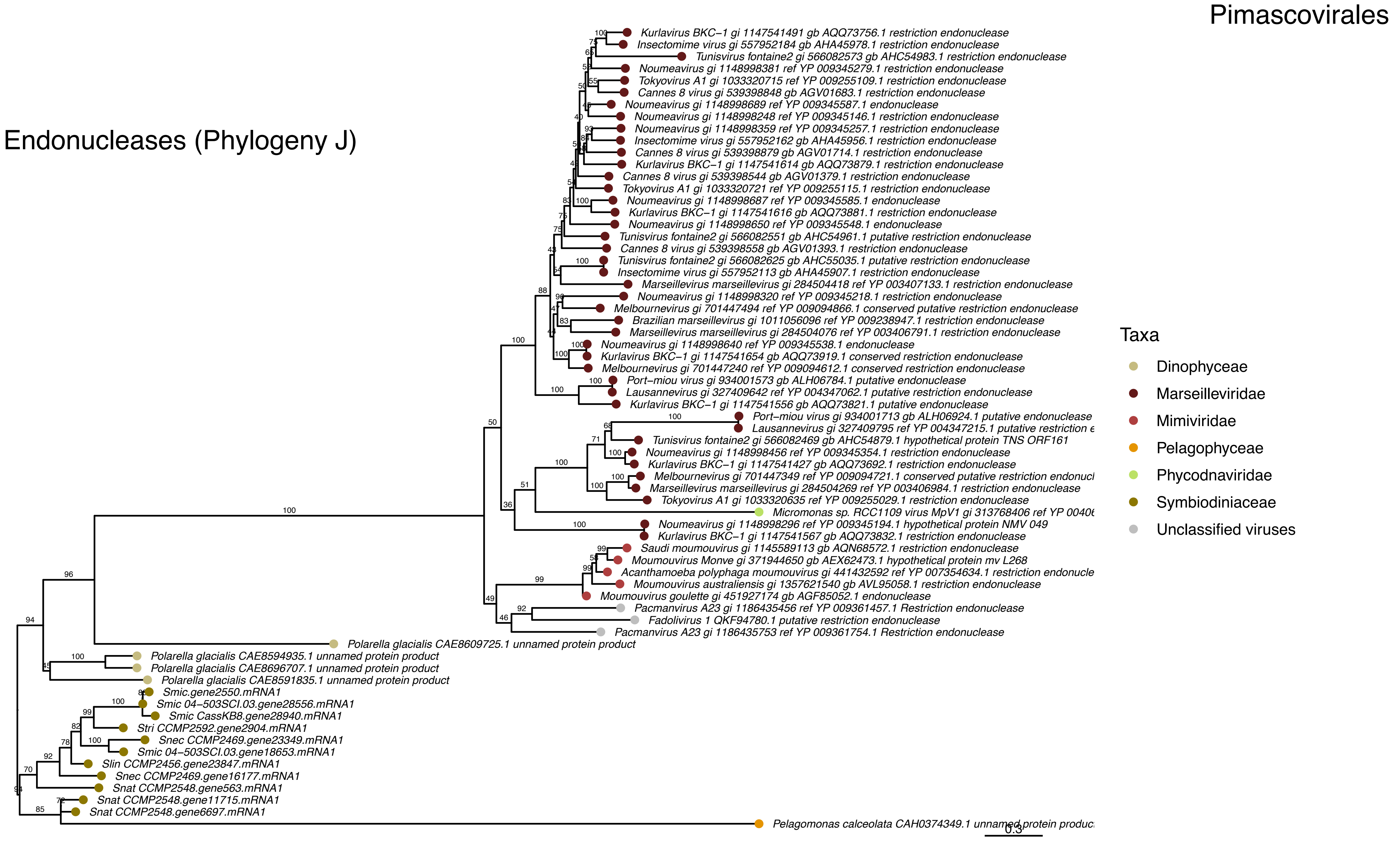
Pimascovirales
Endonucleases (Phylogeny J)
Kurlavirus BKC–1 gi 1147541491 gb AQQ73756.1 restriction endonuclease
Insectomime virus gi 557952184 gb AHA45978.1 restriction endonuclease
Tunisvirus fontaine2 gi 566082573 gb AHC54983.1 restriction endonuclease
Noumeavirus gi 1148998381 ref YP 009345279.1 restriction endonuclease
Tokyovirus A1 gi 1033320715 ref YP 009255109.1 restriction endonuclease
Cannes 8 virus gi 539398848 gb AGV01683.1 restriction endonuclease
Noumeavirus gi 1148998689 ref YP 009345587.1 endonuclease
Noumeavirus gi 1148998248 ref YP 009345146.1 restriction endonuclease
Noumeavirus gi 1148998359 ref YP 009345257.1 restriction endonuclease
Insectomime virus gi 557952162 gb AHA45956.1 restriction endonuclease
Cannes 8 virus gi 539398879 gb AGV01714.1 restriction endonuclease
Kurlavirus BKC–1 gi 1147541614 gb AQQ73879.1 restriction endonuclease
Cannes 8 virus gi 539398544 gb AGV01379.1 restriction endonuclease
Tokyovirus A1 gi 1033320721 ref YP 009255115.1 restriction endonuclease
Noumeavirus gi 1148998687 ref YP 009345585.1 endonuclease
Kurlavirus BKC–1 gi 1147541616 gb AQQ73881.1 restriction endonuclease
Noumeavirus gi 1148998650 ref YP 009345548.1 endonuclease
Tunisvirus fontaine2 gi 566082551 gb AHC54961.1 putative restriction endonuclease
Cannes 8 virus gi 539398558 gb AGV01393.1 restriction endonuclease
Tunisvirus fontaine2 gi 566082625 gb AHC55035.1 putative restriction endonuclease
Insectomime virus gi 557952113 gb AHA45907.1 restriction endonuclease
Marseillevirus marseillevirus gi 284504418 ref YP 003407133.1 restriction endonuclease
Noumeavirus gi 1148998320 ref YP 009345218.1 restriction endonuclease
Melbournevirus gi 701447494 ref YP 009094866.1 conserved putative restriction endonuclease
Brazilian marseillevirus gi 1011056096 ref YP 009238947.1 restriction endonuclease
Marseillevirus marseillevirus gi 284504076 ref YP 003406791.1 restriction endonuclease
Noumeavirus gi 1148998640 ref YP 009345538.1 endonuclease
Kurlavirus BKC–1 gi 1147541654 gb AQQ73919.1 conserved restriction endonuclease
Melbournevirus gi 701447240 ref YP 009094612.1 conserved restriction endonuclease
Port–miou virus gi 934001573 gb ALH06784.1 putative endonuclease
Lausannevirus gi 327409642 ref YP 004347062.1 putative restriction endonuclease
Kurlavirus BKC–1 gi 1147541556 gb AQQ73821.1 putative endonuclease
Port–miou virus gi 934001713 gb ALH06924.1 putative endonuclease
Lausannevirus gi 327409795 ref YP 004347215.1 putative restriction e
Tunisvirus fontaine2 gi 566082469 gb AHC54879.1 hypothetical protein TNS ORF161
Noumeavirus gi 1148998456 ref YP 009345354.1 restriction endonuclease
Kurlavirus BKC–1 gi 1147541427 gb AQQ73692.1 restriction endonuclease
Melbournevirus gi 701447349 ref YP 009094721.1 conserved putative restriction endonucl
Marseillevirus marseillevirus gi 284504269 ref YP 003406984.1 restriction endonuclease
Tokyovirus A1 gi 1033320635 ref YP 009255029.1 restriction endonuclease
Micromonas sp. RCC1109 virus MpV1 gi 313768406 ref YP 0040
Noumeavirus gi 1148998296 ref YP 009345194.1 hypothetical protein NMV 049
Kurlavirus BKC–1 gi 1147541567 gb AQQ73832.1 restriction endonuclease
Saudi moumouvirus gi 1145589113 gb AQN68572.1 restriction endonuclease
Moumouvirus Monve gi 371944650 gb AEX62473.1 hypothetical protein mv L268
Acanthamoeba polyphaga moumouvirus gi 441432592 ref YP 007354634.1 restriction endonucle
Moumouvirus australiensis gi 1357621540 gb AVL95058.1 restriction endonuclease
Moumouvirus goulette gi 451927174 gb AGF85052.1 endonuclease
Pacmanvirus A23 gi 1186435456 ref YP 009361457.1 Restriction endonuclease
Fadolivirus 1 QKF94780.1 putative restriction endonuclease
Pacmanvirus A23 gi 1186435753 ref YP 009361754.1 Restriction endonuclease
Polarella glacialis CAE8609725.1 unnamed protein product
Polarella glacialis CAE8594935.1 unnamed protein product
Polarella glacialis CAE8696707.1 unnamed protein product
Polarella glacialis CAE8591835.1 unnamed protein product
Smic.gene2550.mRNA1
Smic 04–503SCI.03.gene28556.mRNA1
Smic CassKB8.gene28940.mRNA1
Stri CCMP2592.gene2904.mRNA1
Snec CCMP2469.gene23349.mRNA1
Smic 04–503SCI.03.gene18653.mRNA1
Slin CCMP2456.gene23847.mRNA1
Snec CCMP2469.gene16177.mRNA1
Snat CCMP2548.gene563.mRNA1
Snat CCMP2548.gene11715.mRNA1
Snat CCMP2548.gene6697.mRNA1
Pelagomonas calceolata CAH0374349.1 unnamed protein produc
0.3
Taxa
Dinophyceae
Marseilleviridae
Mimiviridae
Pelagophyceae
Phycodnaviridae
Symbiodiniaceae
Unclassified viruses

### Mononegavirales

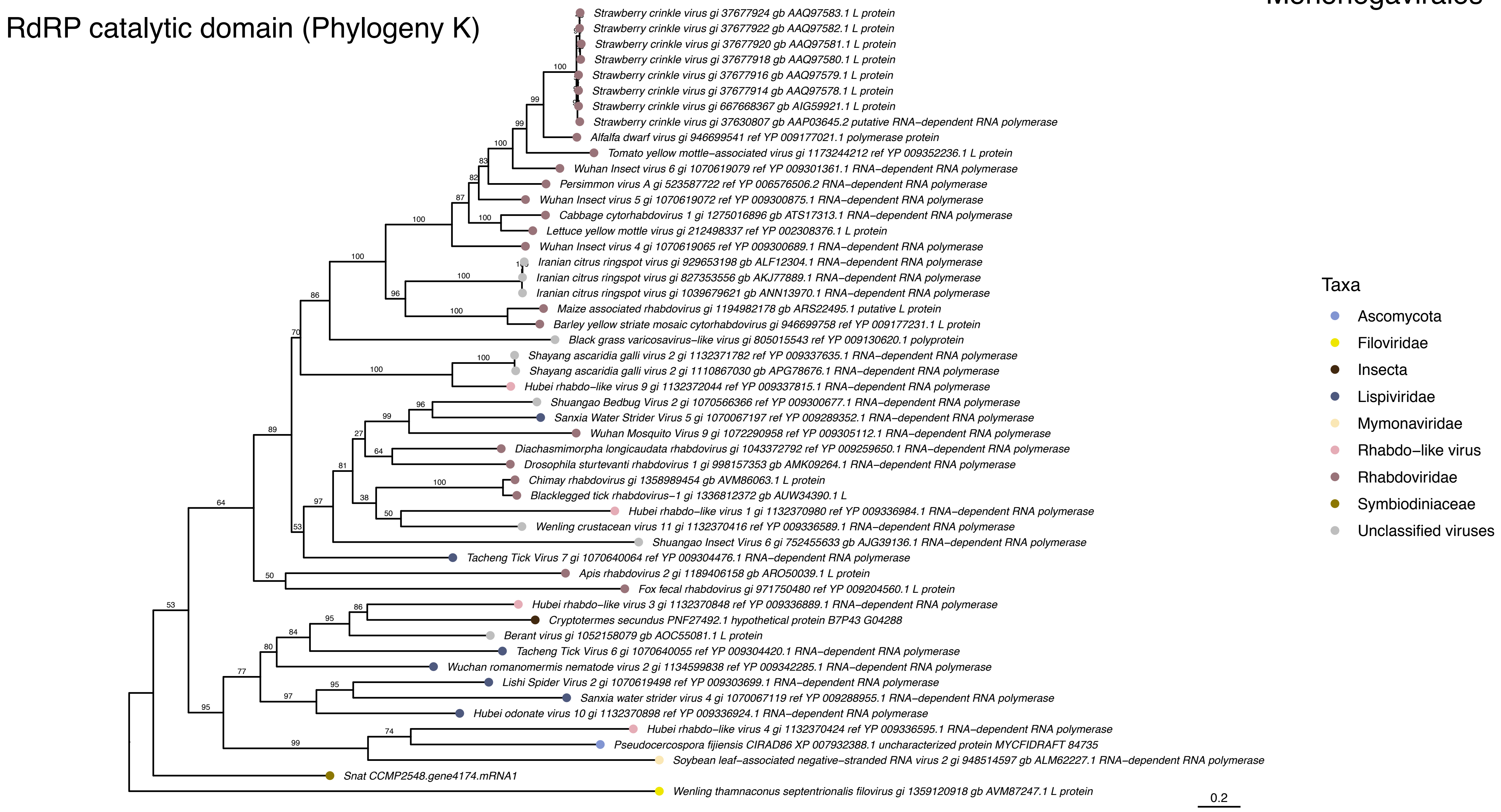


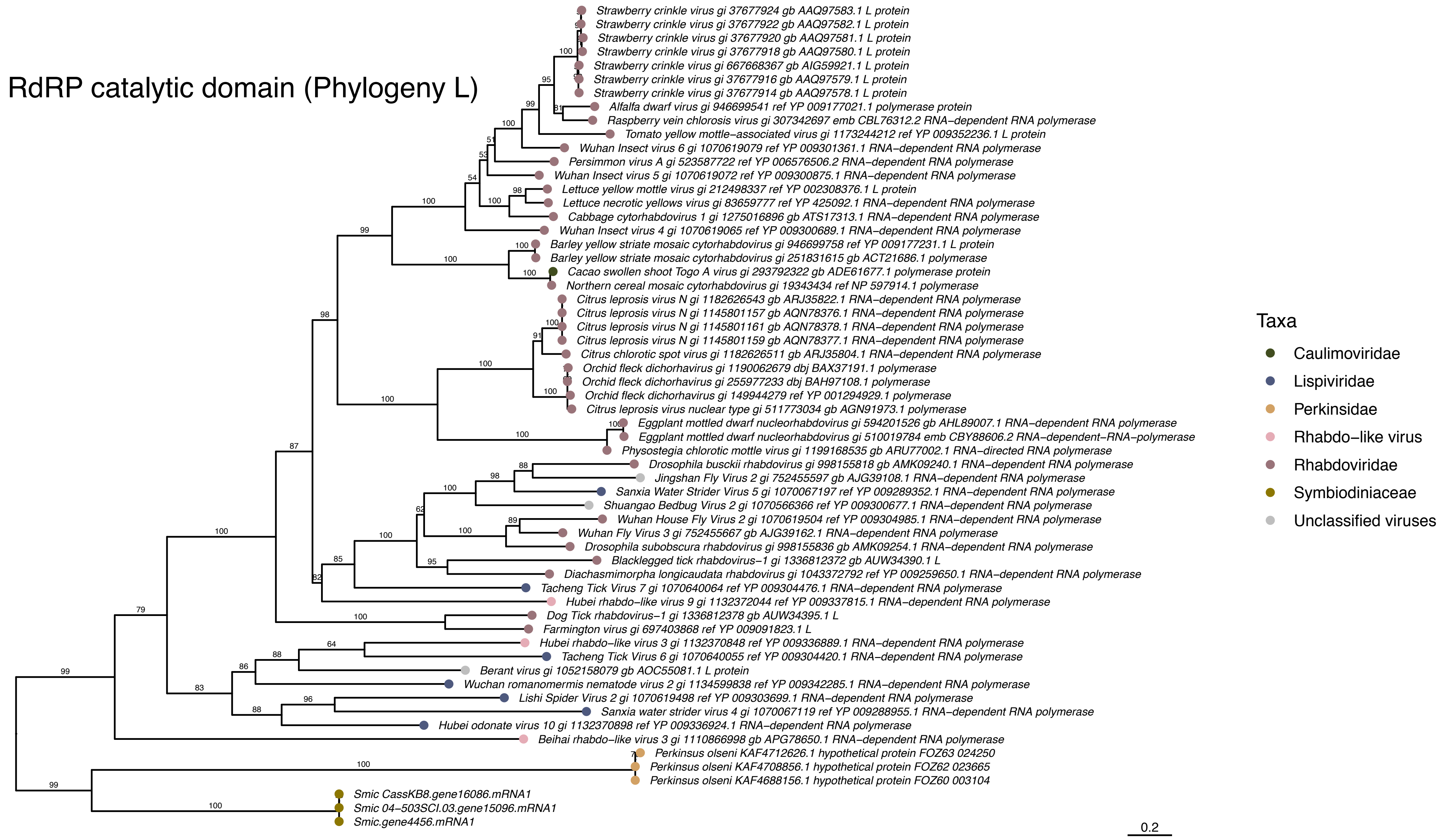
