## Supplementary Fig. S1 for "Multiple waves of viral invasions in Symbiodiniaceae algal genomes"

**A**

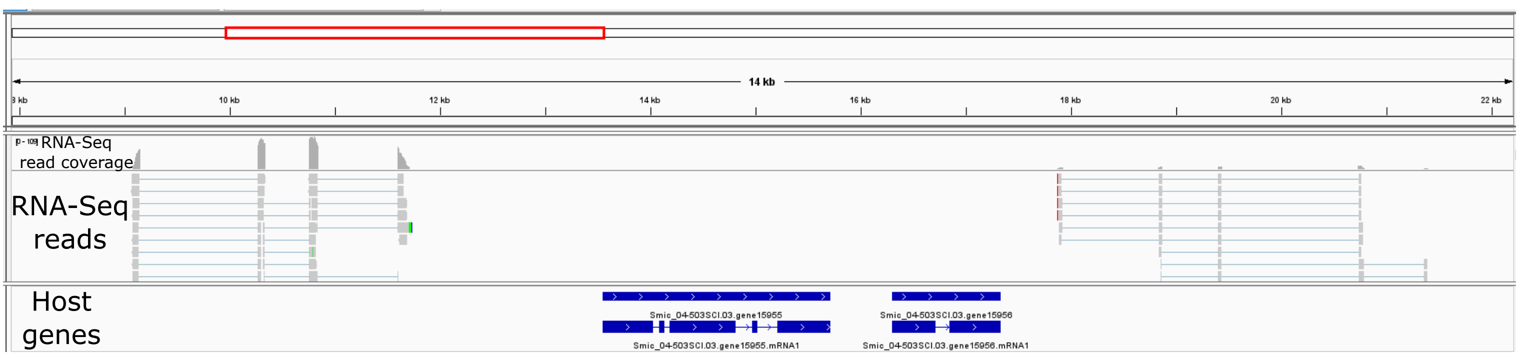


**B**

```
         Query: YP_009337004.1 RNA-dependent RNA polymerase [Beihai weivirus-like virus 5]
        Target: Smic_04-503SCI.03.scaffold3501:13547-15709:+ (Smic_04-503SCI.03.gene15955)
         Model: protein2genome:local
     Raw score: 545
   Query range: 357 -> 767    (Length:820aa; 50% query coverage)
  Target range: 2639 -> 3938

  358 : AspLysArgCysGlyValPheAlaLysHisArgIleGluGluTrpAlaIleAlaHisPheAs :  378
        :!!::!!:!! !  !|||||||::!|||.!.!:!|||!!:!       !:!:::!   ||| !
        AsnArgLysTyrHisValPheSerLysGluGlnIleAspAlaGluIleValSerIlePheHi
 2640 : AACCGGAAGTATCACGTCTTCAGCAAAGAGCAGATTGATGCTGAAATTGTGAGCATATTCCA : 2700

  379 : pLeuGluGluCysLysSerGlyLysTrpSerIleGluArgPheArgGlySerLeuGluAsnL :  399
        !:!!  !:!!! !   |||  !|||||||!!!   :!!|||||||..!..!! !!!!.!.:!!|
        sValProGlnPheAlaSerLysLysTrpThrGlnLysArgPheGluAsnMetPheAsnHisL
 2701 : TGTGCCACAATTCGCCTCCAAGAAATGGACACAGAAAAGATTTGAGAACATGTTCAATCATC : 2763

  400 : euTyrAlaLysGluHisProThrPheSerPheLysAlaAspValLysTyrGlu<-><-><-> :  417
        ||  !  !:!!! ! !!|||  !||||!.!  !||||||| ! :!!|||   |||
        euLeuLysGlnValAspProArgPheLysSerLysAlaLysIleLysLeuGluAlaMetGly
 2764 : TCCTCAAACAGGTGGATCCACGATTCAAGAGCAAGGCCAAAATAAAGCTAGAAGCAATGGGC : 2823

  418 : CysMetProGluGly<-><->LysAlaProArgMetLeuIleAlaAspGlyAspGluGlyGl :  435
        |||!  |||!!:|||      ||| !!|||||||:!!|||||||||||||||||||||||||||
        CysLysProAspGlySerProLysProProArgLeuLeuIleAlaAspGlyAspGluGlyGl
 2824 : TGCAAACCGGATGGCTCGCCAAAGCCACCCCGTCTGCTTATAGCAGACGGGGATGAAGGGCA : 2886

  436 : nLeuMetAlaLeuAlaValValLysCysPheGluGluLeuLeuPheSerHisPheGluThrL :  456
        |:!!|||:!!|||  !! !:!!  !  !||||||:!!||||||||| !! ! |||.!.!:!!
        nIleMetSerLeuLeuAspIleAlaIlePheGluLysLeuLeuPheArgLysPheHisSerA
 2887 : AATCATGTCGCTCCTCGACATAGCAATTTTTGAGAAATTGCTCTTCCGGAAGTTCCACAGTA : 2949

  457 : ysSerIleLysHisLeuAlaLysArgAspAlaIleAspArgValLeuLysGluLeuArgAla :  476
        :!|||||||||  !! !:!!!:!|||.!.!.!:!:.!.  !|||:!!:!! ! |||||| !!
        rgSerIleLysGlyArgSerArgArgGlnValLeuGlnAspValValGluTyrLeuArgPro
 2950 : GGAGCATAAAAGGCCGTTCAAGGAGACAAGTGCTGCAGGACGTGGTCGAGTACTTACGTCCC : 3009

  477 : ProGlyAlaLys  >>>> Target Intron 1 >>>>  AlaValGluGlyAspGlySer :  487
          !  !  ! !              33 bp              !|||||||||| !!||||||
        SerLysLysHis+-                          -+MetValGluGlyHisGlySer
 3010 : AGCAAGAAACATga........................cgATGGTGGAAGGACATGGTTCC : 3075

  488 : AlaTrpAspThrThrCysAsnValLeuIleArgGlyLeuValGluAsnProValLeuArgHi :  508
        |||||||||     !||||::  !  !:!!||||! !:!!|||||||||||||||||||||||||
        AlaTrpAsp---CysCysSerLysGluLeuArgAspMetValGluAsnProValLeuArgHi
 3076 : GCTTGGGAC---TGCTGCTCGAAAGAGCTTAGAGACATGGTAGAAAATCCGGTACTTCGACA : 3135

  509 : sIleThrThrValLeuCysAsnPheGlyValIleProSerThrTrpMetGluGluHisGlnA :  529
        ||||.!!|||  !|||   :!!!:!  !:!!:!!||| !!  !|||  !:!!||||||  !|
        sIleAlaThrHisLeuMetAspTyrTyrLeuValProProGlnTrpGluGlnGluHisAlaA
 3136 : CATCGCGACACATCTGATGGATTACTACCTCGTCCCGCCTCAATGGGAGCAAGAACACGCGA : 3198

  530 : rgAlaCysGluLysLysThrLeuArgLeuPhePheSerAsnLysPheGluThrMetSerThr :  549
        ||.!!  !     ! !   !! !!.!|||.!!||| !!:!!|||.!.  !|||   ! !..!
        rgThrAsnThrAlaAspArgTyrAsnLeuIlePheArgAspLysLeuMetThrTyrPheVal
 3199 : GGACCAACACCGCGGACCGCTACAACCTCATCTTCCGGGACAAGCTGATGACCTACTTTGTA : 3258

  550 : SerIleAspAlaIleArgArgSerGlyHisArgGlyThrSerCysLeuAsnTrpTrpIleAs :  570
        ..!      ! !.!! !|||||||||||||||||||||||||||||||||||||||||:!!||
        Glu***LysGlyThrArgArgSerGlyHisArgGlyThrSerCysLeuAsnTrpTrpValAs
 3259 : GAGTAGAAAGGAACACGTCGATCAGGCCATAGAGGTACGTCTTGCTTGAACTGGTGGGTCAA : 3321

  571 : nPheValLeuTrpValSerSerValPheLysGluProGluArgPheLeuAspValAlaValA :  591
        |||||||||||||  !:!!||||||! !!  .!.|||  !  !!!.||| !!!.!  !  !
        nPheValLeuTrpSerAlaSerValSerIleAsnProTrpValLeuLeuTyrAlaLysGlnA
 3322 : CTTCGTGTTGTGGAGCGCATCAGTATCCATCAACCCTTGGGTATTGCTCTATGCCAAACAGG : 3384

  592 : rgAsnGlyThrAspLeuThrGlyArgSerArgTrpTrpAsnGlyCysPheGluGlyAspAsp :  611
         !  ! !!! !! !:!!!.!|||!:!!!!! !|||   !..  !  !.!.||||||||||||
        spPheCysLysGlyIleAsnGlyLysArgLeuTrpIleArgIleValLeuGluGlyAspAsp
 3385 : ACTTTTGCAAGGGTATAAATGGGAAGAGGCTTTGGATCAGGATCGTCCTGGAGGGAGACGAT : 3444

  612 : SerLeu<-><->CysThrMetArgProProMetValGluGlyAspAlaLeuCysGlnValPh :  630
        ||||||        !  !   |||! !  !     !||||! !||| !!  !   :!!  !||
        SerLeuGluGlyIleCysArgArgLeuGlyAspProGluGluAspProArgThrLysTyrPh
 3445 : TCCCTGGAGGGGATCTGCCGCAGGCTCGGCGATCCTGAAGAGGACCCGAGAACAAAGTACTT : 3507

  631 : eLeuAlaPheTrpLysSerAlaGlyPheAsnMetLysIleValPheCysLysThrArg<->< :  650
        ||||! !!:!||||||| !!  !||||||:!!|||||||||  !   |||  !..!|||
        eLeuAspTyrTrpLysArgGlnGlyPheAspMetLysIleArgGlnCysGlyValArgProA
 3508 : TCTCGACTATTGGAAACGCCAGGGTTTCGATATGAAAATCCGCCAATGTGGGGTTCGTCCAG : 3570

  651 : -><-><-><-><->AlaThrPheValGlyTrpHisValGlyCysThrAspGly<->GluLeu :  664
                      |||  !||||:!!|||   ||| !!  !     !|||  !   !!:|||
        spThrLysPro***Ala***PheIleGlyThrHisPheMetLeuAspAspHisLeuAspLeu
 3571 : ACACGAAGCCATAGGCCTAGTTCATTGGGACTCATTTCATGCTAGATGACCACCTGGATCTC : 3630

  665 : AsnAsnPheArgCysProGluLeuProArgAlaLeuAlaAsnSerGlyValSerValSerPr :  685
        .!...!  !  !   |||!!:|||||||||  !|||.!!||||!:! !!   |||..!!!!||
        GluGlyThrPheValProAspLeuProArgAsnLeuThrAsnAsnTrp---SerThrThrPr
 3631 : GAGGGCACCTTTGTGCCCGACCTGCCCAGGAATCTGACGAACAATTGG---TCAACAACCCC : 3690

  686 : oGluAlaIleLysAlaAlaLysAspMetAsnArgSerAlaValAsnValLeuAlaAlaAlaS :  706
        |! !  !|||:!!!.!  !:!!.!...!!!.  !  !  !||||!..  !  !::!||||||:
        oGlyMetIleGlnValTyrGluGlnGlnLysPheHisLeuValArgGlnAsnSerAlaAlaA
 3691 : GGGAATGATACAAGTGTACGAACAGCAGAAATTCCACCTGGTGAGGCAAAACAGCGCAGCAG : 3753

  707 : erAlaLeuAlaArgAlaSerAspPheAlaGlyIleLeuProSerValSerValLysTyrMet :  726
        !!|||||||||||||||! !||||:!|||||||||||||||! !:!!|||:!!|||||||:!:
        laAlaLeuAlaArgAlaLeuAspTyrAlaGlyIleLeuProMetLeuSerMetLysTyrVal
 3754 : CAGCACTTGCACGTGCTTTGGATTATGCAGGCATACTACCGATGCTGTCGATGAAATATGTC : 3813

  727 : AspPheAlaGluSerValSerArgThrAspPheSerAspArgGluMetSerIleArgAlaPh :  747
        .!.!:!|||   ..!  !! !  !  !||||||  !:!!..!!!::!:|||  !! !|||
        GlnTyrAlaIleGlnCysTyrAlaGlyAspPheHisAsnGluAspLeuSerTyrLeuAlaTh
 3814 : CAATATGCCATTCAATGCTACGCTGGTGACTTCCACAATGAAGATCTCTCGTATCTTGCCAC : 3876

  748 : eGlyGluAspGlyPheSerAlaAsnAlaValArgThrGlnIleMetGluArgAsnIleGly :  767
         ||||||!  |||  !!!!:!!:!!|||||||! !:!!:!!:!!   ..!! !|||  !|||
        rGlyGluAlaGlyThrThrSerHisAlaValIleSerLysValProArgLeuAsnGlyGly
 3877 : AGGTGAAGCAGGTACCACCTCACACGCTGTCATTTCCAAAGTACCAAGGCTGAATGGTGGG : 3938
```

**C**

```
         Query: AOG17586.1 putative major capsid protein [Symbiodinium +ssRNA virus TR74740 c13_g1_i1]
        Target: Smic_04-503SCI.03.scaffold3501:16298-17334:+ (Smic_04-503SCI.03.gene15956)
         Model: protein2genome:local
     Raw score: 167
   Query range: 32 -> 146    (Length:358aa; 32% query coverage)
  Target range: 2083 -> 2428

   33 : ArgArgSerArgAlaArgProAlaAsnGlyGlySerGlnLeuMetGlyIleLysGlnGlyVa :   53
        !:!||||!:!! !  !|||  !  !!.!  !..!..!||||||:!:|||||||!:!||||||||
        LysArgAsnThrMetArgLysArgThrLysSerAspGlnLeuValGlyIleArgGlnGlyVa
 2084 : AAGCGCAACACGATGCGCAAGAGAACTAAGTCTGACCAACTTGTTGGGATTAGACAAGGAGT : 2144

   54 : lGlyAlaIleThrArgGlyProPheGlySerAsnValAlaTyrGlyProHisCysPheAsnA :   74
        |||||||  ! !!!:! !!  !!:!||||||||!:!! !  !!   ..!  !!.!..! !!:!!|
        lGlyAlaAlaProLysArgGlyTyrGlySerSerGlyLysSerSerLeuArgAlaValAspA
 2145 : CGGAGCCGCCCCAAAACGGGGATACGGTTCAAGTGGCAAGTCATCATTGCGCGCCGTCGACG : 2207

   75 : laPheAsnTyrCysHisMetProLeuProArgAlaIleGlyProTyrThrValIleArgThr :   94
        |||||:!!  !||||||:!!||||||||||||||||:!!|||   ! ! |||! !|||||||||
        laPheAspLeuCysHisLeuProLeuProArgAlaValGlyGlyLysThrGlyIleArgThr
 2208 : CTTTTGATCTCTGCCATTTGCCTTTACCCAGGGCAGTCGGTGGTAAAACTGGCATACGGACA : 2267

   95 : ThrArgValIleLysSerAsnLeuGlnLeuMetAsnPheGlyThr<->MetTyrAsnGluAr :  114
        |||  !||||||!  |||:!!  !! !|||   ! !|||||||||   |||   ::!!!:!
        ThrValValIleThrSerAspAspProLeuSerIlePheGlyThrMetMetValSerAspTh
 2268 : ACCGTTGTTATCACCAGTGATGATCCATTGTCCATCTTCGGCACGATGATGGTGTCTGACAC : 2330

  115 : gGlnThrAlaPheAsnSerSerThrTrpSerAsnValCysAlaTrpGlyThrAsnAsnLeuA :  135
        !   .!!  !!!..!.:!!!!!  !||||||.!.:!!  !||||!::..!! !   :!!  !!
        rValAlaGlnLeuGlnAlaArgGluTrpSerGlnLeuIleAlaTyrSerMetProAspAlaG
 2331 : TGTCGCACAATTGCAGGCCAGAGAGTGGAGTCAGTTGATTGCCTACTCCATGCCAGATGCTG : 2393

  136 : laAsnProMetAsnGlyAlaAlaAsnAlaThrArg :  146
        .!.!.! !!!:!.!|||.!!  !  !!.!||||!:!
        lyGlnLeuIleThrGlyThrAsnAlaValThrLys
 2394 : GCCAGCTCATCACTGGGACCAACGCTGTCACTAAG : 2428
```
