## Supplementary Fig. S2 for "Multiple waves of viral invasions in Symbiodiniaceae algal genomes"

**A**

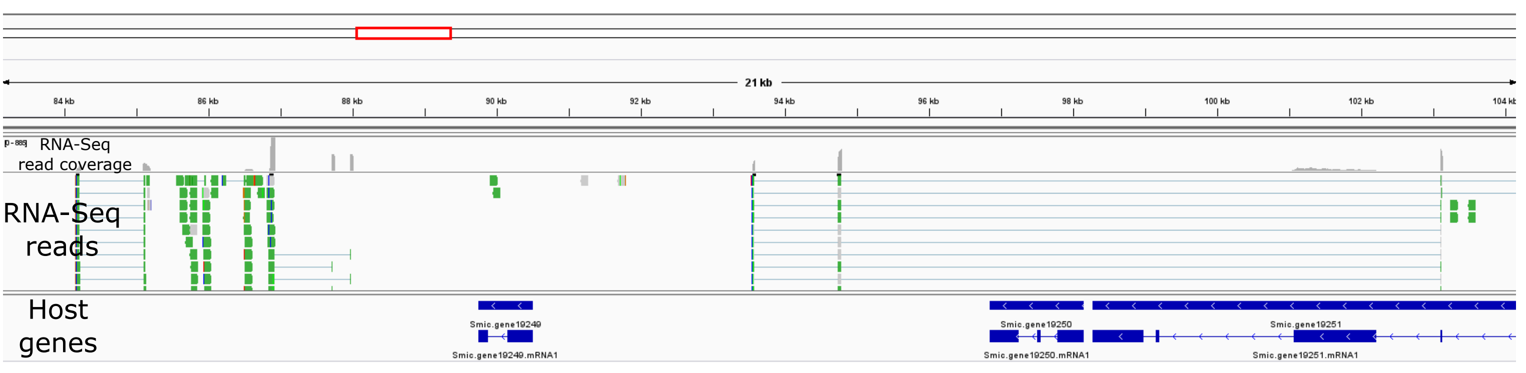


**B**

```
         Query: AOG17586.1 putative major capsid protein [Symbiodinium +ssRNA virus TR74740 c13_g1_i1]
        Target: Smic.scaffold792:89753-90508:- (Smic.gene19249)
         Model: protein2genome:local
     Raw score: 678
   Query range: 28 -> 338    (Length:358aa; 87% query coverage)
  Target range: 1404 -> 2345

   29 : GlnAspSerArgArgArgSerArgAlaArgProAlaAsnGlyGlySerGlnLeuMetGlyIl :   49
        ::!.!...!!:!||||.!:!!|||  !!:! !!!.!:!!  !..!||||||||||:!:|||||
        ArgGlnAspLysArgAsnAlaArgArgLysAlaGlyAspGlnSerSerGlnLeuValGlyIl
 1405 : AGACAAGACAAGCGCAATGCAAGGAGAAAGGCAGGCGATCAATCTTCACAGCTTGTTGGGAT : 1465

   50 : eLysGlnGlyValGlyAlaIleThrArgGlyProPheGlySerAsn<-><->ValAlaTyrG :   68
        |||||||||||!.||||  !  !..!! !  !||||||||||||||!:!      |||:!!||||!
        eLysGlnGlyAlaGlyLysSerValProLysProPheGlySerSerThrArgValSerTyrA
 1466 : TAAGCAGGGAGCAGGCAAGTCCGTCCCAAAACCCTTTGGGTCAAGCACCCGTGTGTCCTATG : 1528

   69 : lyProHisCysPheAsnAlaPheAsnTyrCysHisMetProLeuProArgAlaIleGlyPro :   88
        .! !!!!. !!||||:!!||||||||:!!   ! !||||:!:||||||||||||||||:!:||||||
        laSerGlnSerPheAspAlaPheHisProSerHisLeuProLeuProArgAlaValGlyPro
 1529 : CCTCACAGAGTTTTGATGCCTTCCATCCATCTCATCTACCCTTGCCACGTGCAGTGGGTCCC : 1588

   89 : TyrThrValIleArgThrThrArgValIleLysSerAsnLeuGlnLeuMetAsnPheGlyTh :  109
        ||||! !  !:!:||||||||||! !||||||...!!!!..  !:!!|||:!:! !|||||| !
        TyrIleProValArgThrThrThrValIleSerThrThrAspLysLeuValIlePheGlyPr
 1589 : TATATTCCTGTGCGCACCACTACCGTCATTTCCACAACAGACAAATTGGTCATCTTCGGGCC : 1651

  110 : rMetTyr<->AsnGluArgGlnThrAlaPheAsnSerSerThrTrpSerAsnValCysAlaT :  129
        !|||      ::!!!:! !.!.|||  ! !!   :!!!!!!.!||||||||!!.!.!  !||||!
        oMetMetSerSerAspProAspThrGlnValAlaAlaArgAsnTrpSerLysAlaIleAlaT
 1652 : AATGATGTCTTCTGACCCAGATACACAGGTTGCAGCAAGGAATTGGAGCAAGGCGATTGCTT : 1714

  130 : rpGlyThrAsnAsnLeuAlaAsnProMetAsnGlyAlaAlaAsnAlaThrArgPheValPhe :  149
        ::..!! !  !:!!:!!.!!.!. !!!!:!..!.!.!!! !...!.!  !|||  !!.!.!.
        yrSerMetProAspIleThrGlnAlaIleThrAlaThrAspGlyValTyrArgHisAlaMet
 1715 : ATTCCATGCCTGACATCACGCAAGCTATCACAGCAACCGACGGGGTTTACAGGCATGCAATG : 1774

  150 : AspThrMetSerSerAlaSerTrpAspPheAlaThrCysCysProAlaAlaPheSerIleGl :  170
        |||:!!:!:!!!|||   .!!!:::!!  !::!||||     |||||||||||||||:!:||
        AspSerLeuThrSer---GlyPheAsnGlySerThrLeuThrProAlaAlaPheSerValGl
 1775 : GACTCACTTACTTCT---GGCTTTAACGGCAGCACATTGACACCAGCAGCATTCTCCGTGCA : 1834

  171 : nValMetAsnProAsnAlaLeuGlnThrThrSerGlyIleValTyrGlnGlyArgPheArgT :  191
        |:!!||||||||||||||||||||||||||||:!!|||!!:..!|||..!||||||||||!.!|
        nIleMetAsnProAsnAlaLeuGlnThrThrAlaGlyMetThrTyrMetGlyArgPheHisT
 1835 : AATAATGAATCCAAACGCTTTGCAGACCACTGCTGGGATGACTTACATGGGTCGTTTCCACA : 1897

  192 : hrValProGlyLeuSer-----ThrLeuThrThrGlyLeuAlaPheAlaAspAsnTrpLeuS :  210
        |||||||||! !!!!:!!#####:!!  !! !|||||||! !||||||||||||||!:::!!|
        hrValProGluPheAla#####SerSerIleThrGlyGlnAlaPheAlaAspAsnPheValS
 1898 : CGGTCCCGGAGTTTGCTTTGGGTCAAGCATTACAGGCCAAGCATTTGCCGACAACTTTGTTT : 1959

  211 : erTyrAsnAlaProArgLeuThrSerAlaAlaLysLeuAlaPheArgGlyThrHisIleAsp :  230
        ||||||||||||||||||||  !:!!||||!.!|||||||||.!.||||||..!!!.||||||
        erTyrAsnAlaProArgLeuCysAlaAlaGlyLysLeuAlaMetArgGlyValGlnIleAsp
 1960 : CTTACAATGCACCGCGGCTTTGTGCTGCTGGCAAGTTGGCAATGCGTGGGGTTCAGATTGAC : 2019

  231 : AlaValProPheAsnMetSerLysValSerAspPheGlyGluLeuMetLysTyrValAspAs :  251
        |||:!!||||||||||||||||||:!!:!!|||||| !!...|||    ! ! !  !...
        AlaIleProPheAsnMetSerLysLeuAlaAspPheCysArgLeuAspAspSerGlnSerIl
 2020 : GCGATACCTTTCAACATGAGCAAACTCGCTGATTTCTGTCGCTTGGACGACTCTCAGTCGAT : 2082

  252 : pSerAlaValGlnIleThrAlaAspGluSerLeuProAlaGlyPheAlaProIleCysIleT :  272
        !..!.!!! !  !:!::!!!.!|||... !!!!!  !! !||||||||||||||||||  !!
        eGluThrAspThrLeuSerValAspSerProPhePheAspGlyPheAlaProIleCysCysT
 2083 : TGAGACCGACACGCTGTCTGTCGACTCCCCATTTTTCGACGGTTTCGCTCCCATTTGTTGTT : 2145

  273 : yrAsnProAspGlyIleGluLeuGluTyrLeuValCysCysGluTrpArgValArgPheAsp :  292
        |||||||||!!:.!!||||.!.|||:!!!:!|||||||||  !|||||||||||||||||||||
        yrAsnProGluSerIleAsnLeuGlnPheLeuValCysValGluTrpArgValArgPheAsp
 2146 : ACAATCCCGAGAGCATCAACCTCCAATTTCTCGTTTGCGTTGAGTGGCGTGTCCGGTTTGAC : 2205

  293 : ProSerAsnProAlaGlnAlaAlaHisValHisHisProIleAlaSerAspTrpSerTrpAs :  313
        ||||!!||||||||||||||!.!||||||  !|||  !|||  !:!!||||!!:   :!!|||:!
        ProThrAsnProAlaGlnGlyAlaHisArgHisPheProProSerSerGluAlaAlaTrpHi
 2206 : CCTACCAATCCTGCACAGGGTGCCCATCGGCACTTTCCTCCATCCAGTGAAGCTGCATGGCA : 2268

  314 : nSerAlaLeuArgAlaLeuGluAlaGlyAlaHisGlyValLysAspIleValGluThrValA :  334
        !!.!|||:!!  !:!!  !   |||  !!.!:!!|||||||||||||||||||||..!||||
        sLysAlaValSerSerAlaPheAlaIleGlyAsnGlyValLysAspIleValGluValValA
 2269 : CAAGGCCGTATCATCTGCATTCGCGATAGGGAATGGTGTCAAGGACATTGTCGAAGTTGTTG : 2331

  335 : laAsnLeuGlyGln :  338
        ||!:!  !||||||
        laSerThrGlyGln
 2332 : CAAGCACAGGCCAG : 2345
```

**C**

```
         Query: YP_009342067.1 RNA-dependent RNA polymerase [Wenzhou weivirus-like virus 1]
        Target: Smic.scaffold792:96851-98156:- (Smic.gene19250)
         Model: protein2genome:local
     Raw score: 600
   Query range: 318 -> 786  (Length:840aa; 56% query coverage)
  Target range: 1839 -> 3286

  319 : ValValGlyValLeuValGlyProCysGlnArgLysProAsnValTyrSerLysThrValGl :  339
        ||||||||||||  !||||||||||  !  !..!:!!|||   :!!|||.!!!!.  !:!!!!
        ValValGlyValAlaValGlyProLeuAlaGluGluProValIleTyrGlyAsnGluIleAs
 1840 : GTCGTGGGCGTTGCAGTTGGACCTCTCGCAGAGGAACCAGTGATTTACGGGAATGAGATCGA : 1900

  340 : uAsnLeuLysAlaAlaIleAsnGluArgIleIleLysLysAlaArgLysProLysLeuCysA :  360
        :|||:!!:!!||||||  !!!.||||||  !:!:!:!     !! !  ! !!  !! !   !
        pAsnIleGlnAlaAlaAlaLysGluArgAlaLeuArgAlaPheProLeuThrValArgProL
 1901 : TAATATCCAAGCTGCAGCAAAAGAGCGCGCATTGAGGGCTTTCCCGTTGACAGTACGACCGA : 1963

  361 : rgAlaAspArgValArgIleSerArgLeuValArgLysAlaMetSerAsnSerGluLysHis :  380
        :!.!!|||||||  !  !  !..!  !|||:!!||||::!  !   ||||!..  !:!!!:!:!!
        ysThrAspArgArg***GlyGluSerLeuLeuArgArgTrpAlaSerArgValGlnArgAsn
 1964 : AGACAGACAGAAGATAGGGAGAGTCATTGCTACGTCGATGGGCTTCCAGGGTCCAAAGAAAC : 2023

  381 : --GlyValPheSerLysSerArgIleGlnGluTrpAlaIleGlnHisPheAsnLeuGluGlu :  400
        ##  !!.!||||||||||..!!:!|||:!!.!.     !    !   !|||:!!!!!..!!!:
        ##HisAlaPheSerLysGluGlnIleGluAsnGluLeuGluTyrIlePheHisPheSerAsp
 2024 : TTCATGCATTTTCAAAGGAGCAAATCGAAAATGAACTCGAGTATATCTTCCACTTCTCAGAC : 2085

  401 : MetLysSerGlnLysTrpSerLeuGluArgPheArgAsnAlaLeuLysAsnLeuTyrGluLy :  421
        :!:||||||:!!|||||||||!!!  !:!!||||||||  !|||.!!!!!:!!.!.|||   ...:!
        LeuLysSerLysLysTrpThrAsnLysArgPheTyrAsnThrPheGluGluLeuLeuArgGl
 2086 : CTCAAATCCAAGAAATGGACGAATAAGAGGTTCTATAACACTTTCGAAGAGCTCCTGCGCGA : 2148

  422 : sAspProValGluPheMetLeuLysAlaGlyIleLysProGluCysMet<-><-><->GluG :  439
        !!    !  !..!|||! !:!!!  |||  !||||||! !|||   |||         ..!!
        uValAspProArgPheLysMetThrAlaLysIleLysLeuGluProMetGlyLysArgSerA
 2149 : AGTGGACCCACGGTTCAAGATGACTGCCAAGATAAAACTTGAACCGATGGGCAAGCGGTCAG : 2211

  440 : luGly<-><->LysAlaProArgMetLeuIleAlaAspGlyAspGluGlyGlnLeuMetAla :  457
        !:|||      ||| !!||||||:!:||||||||||||||||||||||||||||:!!||||||
        spGlyThrProLysProProArgLeuLeuIleAlaAspGlyAspGluGlyGlnIleMetAla
 2212 : ACGGAACACCAAAGCCCCCAAGGCTTCTCATAGCTGACGGCGACGAAGGACAGATCATGGCT : 2271

  458 : LeuAlaValIleLysCysPheGluGluLeuLeuPheLysHisPheGluGluArgSerIleLy :  478
        |||  !  !:!!..!  !||||||..!|||||| ! !:! ! !:::!!:!!|||||!!|||||
        LeuPheCysValSerIlePheGluArgLeuLeuValArgLysTrpLysLysArgThrIleLy
 2272 : CTGTTCTGCGTCTCGATCTTTGAGAGGCTCCTTGTGAGGAAGTGGAAGAAGAGGACGATTAA : 2334

  479 : sHisLeuSerLysHisGluAlaMetGluArgValValArgValLeuLysLysLysGlyAlaA :  499
        |  !! !|||   !..|||||||!!:.!.  !|||:!!  !  !|||:::   ...  !  !!
        sGlyArgSerGlyArgGluAlaIleAsnAspValIleSerTyrLeuArgProSer***LysH
 2335 : GGGCCGCAGCGGCCGGGAGGCCATTAACGATGTGATCTCGTACCTTCGTCCCTCTTAAAAAC : 2397

  500 : rg  >>>> Target Intron 1 >>>>  AlaValGluGlyAspGlySerAlaTrpAspT :  510
        .!             30 bp                  ||||||||||||||||||||||||:
        is+-                          -+------GluGlyAspGlySerAlaTrpAspS
 2398 : ACgg........................tg------GAGGGGGACGGGTCCGCCTGGGACT : 2454

  511 : hrThrCysGlyValLeuValArgSerLeuIleGluAsnProIleLeuPheHisIlePheArg :  530
        !!  !|||..!  !  !||||||..!|||:!:|||||||||:!!|||   ||||||||||! !
        erCysCysSerLysGlyValArgAspLeuValGluAsnProValLeuGlnHisIlePheThr
 2455 : CGTGTTGTTCCAAAGGCGTGCGAGACCTTGTGGAAAACCCGGTCTTACAACACATATTCACA : 2514

  531 : ThrLeuAlaGluPheGlyValIleProGluThrTrpMetAlaGluHisHisAlaAlaCysTh :  551
        ..!|||||||!!:!:!  !||||||||||..!  !||||||||||!  |||!!.  !!.!  !!
        ValLeuAlaAspTyrMetValIleProSerGlnTrpMetAlaAlaHisGlnAsnValAsnMe
 2515 : GTGCTGGCTGACTACATGGTCATTCCATCACAGTGGATGGCAGCCCACCAGAATGTCAACAT : 2577

  552 : rLysGluLysLeuLysLeuPhePheSerAsnLysPheGlnLysValThrValThrIleAspA :  572
        !:!!..!::!||||!  |||.!!||||!!!:!!|||.!.:!!!    !  !||||!.!|||   |
        tGlnSerArgLeuThrLeuIlePheArgAspLysLeuLysThrHisPheValAsnIleArgA
 2578 : GCAATCGCGGTTGACCCTGATCTTCAGGGACAAGCTGAAGACCCACTTTGTCAACATCAGGG : 2640

  573 : laIleArgArgSerGlyHisArgGlyThrSerCysLeuAsnTrpTrpIleAsnPheThrMet :  592
        ||||||||||||||||||||||||||||||||||||||||||||||:!!|||||||..!:!!
        laIleArgArgSerGlyHisArgGlyThrSerCysLeuAsnTrpTrpValAsnPheValLeu
 2641 : CAATAAGGAGGTCTGGACACCGCGGTACCAGTTGCTTAAACTGGTGGGTAAACTTTGTGTTG : 2700

  593 : TrpValSerSerIlePheLysGluProGluArgPheLeuAsp--ProAspValArgLysGly :  612
        ||||!.!!!!||||||! !...!!:|||  !  !.!.|||:!!## !!|||
        TrpAlaCysSerIleSerSerAspProTrpValLeuLeuAsn##AlaAsp------------
 2701 : TGGGCATGCAGCATTTCTTCCGACCCATGGGTTCTACTCAATGTGCCGAC------------ : 2753

  613 : ThrAspLeuThrGluLysGluArgTrpTrpAsnGlyThrPheGluGlyAspAspSerLeuCy :  633
                     !  |||  !||||||||||...|||..!|||||||||||| !!|||||||
        ------------GlyLysTrpArgTrpTrpArgGlyValPheGluGlyAspHisSerLeuTh
 2754 : ------------GGCAAGTGGAGATGGTGGCGGGGGGTGTTCGAAGGCGACCATTCTCTGAC : 2801

  634 : sThrMetCysProProMet<-><->LeuLysAspAspLysMetSerAspIlePheCysGlnP :  652
         ! !! !||||||||||:!!      ! !:!!|||||||!  !  !!!! !  !||||!  :!!|
        rArgThrCysProProLeuGlyProProGluAspAspThrAsnThrValTyrPheLeuGluP
 2802 : GAGAACGTGCCCCCCACTGGGCCCACCAGAAGACGACACCAACACTGTCTATTTTTTGGAGT : 2864

  653 : heTrpSerAspAlaGlyPheAsnMetLysIleValTyrCysAspArgArg<-><-><-><-> :  669
        |||||!.!  !..!|||||||||||||||:!!  ! !!|||! !  !|||
        heTrpLysArgCysGlyPheAsnMetLysLeuArgAsnCysGlyLeuArgProAspThrArg
 2865 : TTTGGAAGCGCTGCGGGTTTAACATGAAGTTAAGGAACTGTGGCTTGAGGCCTGACACTCGC : 2924

  670 : <-><->AlaThrPheThrGlyTrpHisLeuValCys<->GluAsnGlyGluLeuThrAspVa :  686
              |||  !||||! !|||   |||!!!|||  !   !!:|||  !!!:|||  !!  !
        LysLeuAlaGluPheIleGlyThrHisPheValLeuAspAspAsnMetAspLeuAspGlyAs
 2925 : AAGCTTGCGGAGTTCATCGGGACACACTTTGTCCTTGATGACAACATGGATCTGGACGGGGA : 2987

  687 : lArgAlaProGluLeuProArgAlaLeuAlaAsnSerGlyValSerValSerProSerAlaV :  707
        ! !!  !||||!!:|||||||||  !|||  !|||! !   ||||||  !||||||..!  !:
        pTrpLeuProAspLeuProArgAsnLeuArgAsnPhe---ValSerSerSerProGlnMetI
 2988 : TTGGCTACCAGACCTTCCCCGCAACTTGCGTAATTTT---GTCTCTTCCTCACCACAGATGA : 3047

  708 : alGlnAlaAlaLysAspGlyLysMetSerAlaIleLysValIleAlaAlaAlaSerAlaLeu :  727
        !!||||||  !:!!.!.|||:!!  !  !  !:!!!:!  !||||||||||||:!!  !|||
        leGlnAlaPheGluGlnGlyGlnTrpHisLeuValArgGlnIleAlaAlaAlaAlaAsnLeu
 3048 : TCCAAGCCTTTGAACAGGGCCAGTGGCACCTCGTCAGGCAAATTGCAGCGGCAGCAAATTTA : 3107

  728 : AlaArgAlaSerAspPheSerGlyIleLeuProThrValSerAsnLysTyrLeuGlnTyrAl :  748
        :!!||||||||! !||||:!:!!||||!    !||||! !:!!|||.!.|||||||||   |||||
        SerArgAlaIleAspTyrAlaGlyLysAlaProMetLeuSerGlnLysTyrLeuAlaTyrAl
 3108 : TCTCGCGCCATCGATTATGCCGGCAAGGCACCCATGTTGTCGCAGAAATATTTGGCTTACGC : 3170

  749 : aAsnGluCysSerLysSerAspTyrValAspArgGluCysSerIleArgCysPheGlyPheA :  769
        ||||..!  !|||   ..!||||!:!|||:!!  !!!:   |||!!:! !..!  !|||  !
        aAsnSerIleSerCysGluAspPheValAsnAspAspMetSerMetLeuAlaThrGlyAspT
 3171 : GAACTCAATCAGCTGTGAGGATTTTGTGAATGATGACATGTCGATGCTCGCCACTGGCGATA : 3233

  770 : spGlyHisSerAlaLysGluValArgGluArgIleGluGluArgAsnLeuGly :  786
          |||   !!!|||  !  !||||||  !! !|||.!.:!!  !|||! !|||
        hrGlyAlaThrAlaAlaThrValArgMetMetIleHisGlnTyrAsnSerGly
 3234 : CGGGAGCAACAGCAGCAACAGTTCGCATGATGATACATCAGTACAACTCCGGG : 3286
```
