## Supplementary Fig. S3 for "Multiple waves of viral invasions in Symbiodiniaceae algal genomes"

**A**

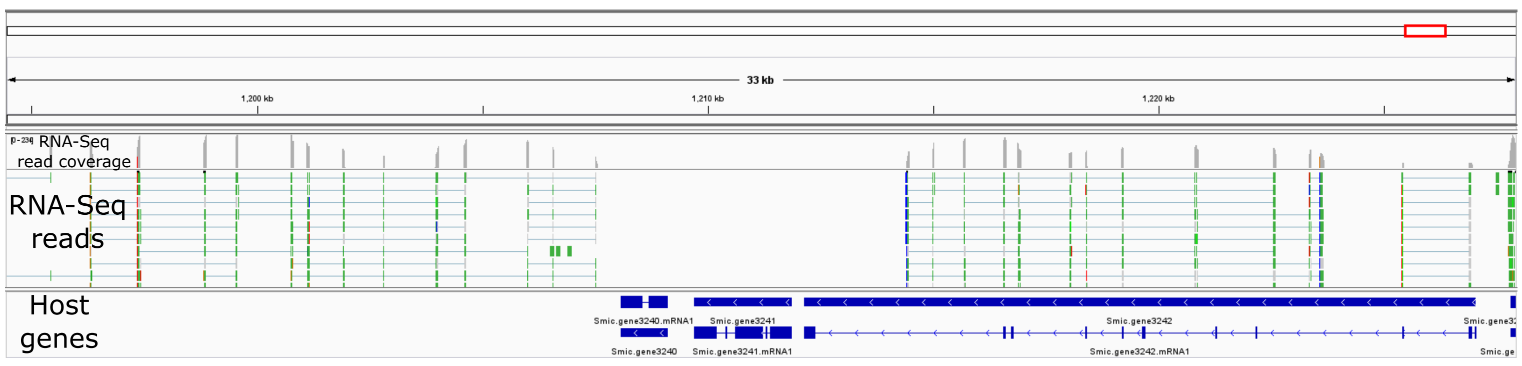


**B**

```
         Query: AOG17586.1 putative major capsid protein [Symbiodinium +ssRNA virus TR74740 c13_g1_i1]
        Target: Smic.scaffold67:1208081-1209117:- (Smic.gene3240)
         Model: protein2genome:local
     Raw score: 167
   Query range: 32 -> 146    (Length:358aa; 32% query coverage)
  Target range: 2084 -> 2429

  33 : ArgArgSerArgAlaArgProAlaAsnGlyGlySerGlnLeuMetGlyIleLysGlnGlyVa :   53
       !:!||||!:!! !  !|||  !  !!.!  !..!..!||||||:!:||||||!:!||||||||
       LysArgAsnThrMetArgLysArgThrLysSerAspGlnLeuValGlyIleArgGlnGlyVa
2085 : AAGCGCAACACGATGCGCAAGAGAACTAAGTCTGACCAACTTGTTGGGATTAGACAAGGAGT : 2145

  54 : lGlyAlaIleThrArgGlyProPheGlySerAsnValAlaTyrGlyProHisCysPheAsnA :   74
       |||||||  ! !!!:! !!  !!:!||||||||!:!! !  !!  ..!  !!.!..! !!:!!|
       lGlyAlaAlaProLysArgGlyTyrGlySerSerGlyLysSerSerLeuArgAlaValAspA
2146 : CGGAGCCGCCCCAAAACGGGGATACGGTTCAAGTGGCAAGTCATCATTGCGCGCCGTCGACG : 2208

  75 : laPheAsnTyrCysHisMetProLeuProArgAlaIleGlyProTyrThrValIleArgThr :   94
       |||||:!!  !||||||:!!|||||||||||||||:!!|||  ! ! |||! !||||||||||
       laPheAspLeuCysHisLeuProLeuProArgAlaValGlyGlyLysThrGlyIleArgThr
2209 : CTTTTGATCTCTGCCATTTGCCTTTACCCAGGGCAGTCGGTGGTAAAACTGGCATACGGACA : 2268

  95 : ThrArgValIleLysSerAsnLeuGlnLeuMetAsnPheGlyThr<->MetTyrAsnGluAr :  114
       |||  !|||||||!  |||:!!  !! !|||   ! !|||||||||   |||   ::!!!:!
       ThrValValIleThrSerAspAspProLeuSerIlePheGlyThrMetMetValSerAspTh
2269 : ACCGTTGTTATCACCAGTGATGATCCATTGTCCATCTTCGGCACGATGATGGTGTCTGACAC : 2331

 115 : gGlnThrAlaPheAsnSerSerThrTrpSerAsnValCysAlaTrpGlyThrAsnAsnLeuA :  135
       !   .!!  !!!..!.:!!!!!  !||||||.!.:!!  !||||!::..!! !   :!!  !!
       rValAlaGlnLeuGlnAlaArgGluTrpSerGlnLeuIleAlaTyrSerMetProAspAlaG
2332 : TGTCGCACAATTGCAGGCCAGAGAGTGGAGTCAGTTGATTGCCTACTCCATGCCAGATGCTG : 2394

 136 : laAsnProMetAsnGlyAlaAlaAsnAlaThrArg :  146
       .!.!.! !!!:!.!|||.!!  !  !!.!||||!:!
       lyGlnLeuIleThrGlyThrAsnAlaValThrLys
2395 : GCCAGCTCATCACTGGGACCAACGCTGTCACTAAG : 2429
```

**C**

```
         Query: YP_009337004.1 RNA-dependent RNA polymerase [Beihai weivirus-like virus 5]
        Target: Smic.scaffold67:1209706-1211868:- (Smic.gene3241)
         Model: protein2genome:local
     Raw score: 545
   Query range: 357 -> 767   (Length:820aa; 50% query coverage)
  Target range: 2640 -> 3939

 358 : AspLysArgCysGlyValPheAlaLysHisArgIleGluGluTrpAlaIleAlaHisPheAs :  378
       :!!::!!:!! !  !||||||::!|||.!.!:!|||||!:!       !:!:::!   ||| !
       AsnArgLysTyrHisValPheSerLysGluGlnIleAspAlaGluIleValSerIlePheHi
2641 : AACCGGAAGTATCACGTCTTCAGCAAAGAGCAGATTGATGCTGAAATTGTGAGCATATTCCA : 2701

 379 : pLeuGluGluCysLysSerGlyLysTrpSerIleGluArgPheArgGlySerLeuGluAsnL :  399
       !:!!  !:!!! !   |||  !|||||||!!   :!!|||||||..!..!! !!!!.!.:!!|
       sValProGlnPheAlaSerLysLysTrpThrGlnLysArgPheGluAsnMetPheAsnHisL
2702 : TGTGCCACAATTCGCCTCCAAGAAATGGACACAGAAAAGATTTGAGAACATGTTCAATCATC : 2764

 400 : euTyrAlaLysGluHisProThrPheSerPheLysAlaAspValLysTyrGlu<-><-><-> :  417
       ||  !  !:!!! ! !!|||  !||||!.!  !||||||| ! :!!|||   |||
       euLeuLysGlnValAspProArgPheLysSerLysAlaLysIleLysLeuGluAlaMetGly
2765 : TCCTCAAACAGGTGGATCCACGATTCAAGAGCAAGGCCAAAATAAAGCTAGAAGCAATGGGC : 2824

 418 : CysMetProGluGly<-><->LysAlaProArgMetLeuIleAlaAspGlyAspGluGlyGl :  435
       |||!  |||!!:|||      ||| !!|||||||:!!|||||||||||||||||||||||||||
       CysLysProAspGlySerProLysProProArgLeuLeuIleAlaAspGlyAspGluGlyGl
2825 : TGCAAACCGGATGGCTCGCCAAAGCCACCCCGTCTGCTTATAGCAGACGGGGATGAAGGGCA : 2887

 436 : nLeuMetAlaLeuAlaValValLysCysPheGluGluLeuLeuPheSerHisPheGluThrL :  456
       |:!!|||:!!|||  !! !:!!  !  !||||||:!!||||||||||| !! ! |||.!.!:!!
       nIleMetSerLeuLeuAspIleAlaIlePheGluLysLeuLeuPheArgLysPheHisSerA
2888 : AATCATGTCGCTCCTCGACATAGCAATTTTTGAGAAATTGCTCTTCCGGAAGTTCCACAGTA : 2950

 457 : ysSerIleLysHisLeuAlaLysArgAspAlaIleAspArgValLeuLysGluLeuArgAla :  476
       :!|||||||||  !! !:!!!:!|||.!.!.!:!:.!.  !|||:!!:!! ! |||||| !!
       rgSerIleLysGlyArgSerArgArgGlnValLeuGlnAspValValGluTyrLeuArgPro
2951 : GGAGCATAAAAGGCCGTTCAAGGAGACAAGTGCTGCAGGACGTGGTCGAGTACTTACGTCCC : 3010

 477 : ProGlyAlaLys  >>>> Target Intron 1 >>>>  AlaValGluGlyAspGlySer :  487
         !  !  ! !               33 bp               !||||||||| !!||||||
       SerLysLysHis+-                          -+MetValGluGlyHisGlySer
3011 : AGCAAGAAACATga.........................cgATGGTGGAAGGACATGGTTCC : 3076

 488 : AlaTrpAspThrThrCysAsnValLeuIleArgGlyLeuValGluAsnProValLeuArgHi :  508
       |||||||||     !||||:::  !  !:!!||||! !:!!||||||||||||||||||||||||
       AlaTrpAsp---CysCysSerLysGluLeuArgAspMetValGluAsnProValLeuArgHi
3077 : GCTTGGGAC---TGCTGCTCGAAAGAGCTTAGAGACATGGTAGAAAATCCGGTACTTCGACA : 3136

 509 : sIleThrThrValLeuCysAsnPheGlyValIleProSerThrTrpMetGluGluHisGlnA :  529
       ||||.!!|||  !|||   :!!!:!  !:!!:!!||| !!  !|||  !:!!|||||||  !|
       sIleAlaThrHisLeuMetAspTyrTyrLeuValProProGlnTrpGluGlnGluHisAlaA
3137 : CATCGCGACACATCTGATGGATTACTACCTCGTCCCGCCTCAATGGGAGCAAGAACACGCGA : 3199

 530 : rgAlaCysGluLysLysThrLeuArgLeuPhePheSerAsnLysPheGluThrMetSerThr :  549
       ||.!!  !     ! !   !! !!.!|||.!!||| !!:!!|||.!.  !|||   ! !..!
       rgThrAsnThrAlaAspArgTyrAsnLeuIlePheArgAspLysLeuMetThrTyrPheVal
3200 : GGACCAACACCGCGGACCGCTACAACCTCATCTTCCGGGACAAGCTGATGACCTACTTTGTA : 3259

 550 : SerIleAspAlaIleArgArgSerGlyHisArgGlyThrSerCysLeuAsnTrpTrpIleAs :  570
       ..!    ! !.!! !|||||||||||||||||||||||||||||||||||||||||:!!||
       Glu***LysGlyThrArgArgSerGlyHisArgGlyThrSerCysLeuAsnTrpTrpValAs
3260 : GAGTAGAAAGGAACACGTCGATCAGGCCATAGAGGTACGTCTTGCTTGAACTGGTGGGTCAA : 3322

 571 : nPheValLeuTrpValSerSerValPheLysGluProGluArgPheLeuAspValAlaValA :  591
       |||||||||||||  !:!!||||||||! !!  .!.|||  !  !!!.||| !!!.!  !  !
       nPheValLeuTrpSerAlaSerValSerIleAsnProTrpValLeuLeuTyrAlaLysGlnA
3323 : CTTCGTGTTGTGGAGCGCATCAGTATCCATCAACCCTTGGGTATTGCTCTATGCCAAACAGG : 3385

 592 : rgAsnGlyThrAspLeuThrGlyArgSerArgTrpTrpAsnGlyCysPheGluGlyAspAsp :  611
        !  ! !!! !! !:!!!.!||||:!!!!! !|||   !..  !  !.!.||||||||||||
       spPheCysLysGlyIleAsnGlyLysArgLeuTrpIleArgIleValLeuGluGlyAspAsp
3386 : ACTTTTGCAAGGGTATAAATGGGAAGAGGCTTTGGATCAGGATCGTCCTGGAGGGAGACGAT : 3445

 612 : SerLeu<-><->CysThrMetArgProProMetValGluGlyAspAlaLeuCysGlnValPh :  630
       ||||||        !  !   |||! !  !     !||||! !||| !!  !   :!!  !||
       SerLeuGluGlyIleCysArgArgLeuGlyAspProGluGluAspProArgThrLysTyrPh
3446 : TCCCTGGAGGGGATCTGCCGCAGGCTCGGCGATCCTGAAGAGGACCCGAGAACAAAGTACTT : 3508

 631 : eLeuAlaPheTrpLysSerAlaGlyPheAsnMetLysIleValPheCysLysThrArg<->< :  650
       ||||! !!:!||||||| !!  !||||||:!!|||||||||  !   |||  !..!|||
       eLeuAspTyrTrpLysArgGlnGlyPheAspMetLysIleArgGlnCysGlyValArgProA
3509 : TCTCGACTATTGGAAACGCCAGGGTTTCGATATGAAAATCCGCCAATGTGGGGTTCGTCCAG : 3571

 651 : -><-><-><-><->AlaThrPheValGlyTrpHisValGlyCysThrAspGly<->GluLeu :  664
                     |||  !||||:!!|||   ||| !!  !     !|||  !   !!:|||
       spThrLysPro***Ala***PheIleGlyThrHisPheMetLeuAspAspHisLeuAspLeu
3572 : ACACGAAGCCATAGGCCTAGTTCATTGGGACTCATTTCATGCTAGATGACCACCTGGATCTC : 3631

 665 : AsnAsnPheArgCysProGluLeuProArgAlaLeuAlaAsnSerGlyValSerValSerPr :  685
       .!...!  !  !   ||||!:|||||||||  !|||.!!|||||!:! !!   |||..!!!!||
       GluGlyThrPheValProAspLeuProArgAsnLeuThrAsnAsnTrp---SerThrThrPr
3632 : GAGGGCACCTTTGTGCCCGACCTGCCCAGGAATCTGACGAACAATTGG---TCAACAACCCC : 3691

 686 : oGluAlaIleLysAlaAlaLysAspMetAsnArgSerAlaValAsnValLeuAlaAlaAlaS :  706
       |! !  !|||:!!!.!  !:!!.!...!!!.  !  !  !||||!..  !  !::!||||||:
       oGlyMetIleGlnValTyrGluGlnGlnLysPheHisLeuValArgGlnAsnSerAlaAlaA
3692 : GGGAATGATACAAGTGTACGAACAGCAGAAATTCCACCTGGTGAGGCAAAACAGCGCAGCAG : 3754

 707 : erAlaLeuAlaArgAlaSerAspPheAlaGlyIleLeuProSerValSerValLysTyrMet :  726
       !!|||||||||||||||! !||||!:!||||||||||||||! !:!!|||:!!|||||||:!:
       laAlaLeuAlaArgAlaLeuAspTyrAlaGlyIleLeuProMetLeuSerMetLysTyrVal
3755 : CAGCACTTGCACGTGCTTTGGATTATGCAGGCATACTACCGATGCTGTCGATGAAATATGTC : 3814

 727 : AspPheAlaGluSerValSerArgThrAspPheSerAspArgGluMetSerIleArgAlaPh :  747
       .!.!:!|||   ..!  !! !  !  !||||||  !:!!..!!!::!:|||  !! !|||
       GlnTyrAlaIleGlnCysTyrAlaGlyAspPheHisAsnGluAspLeuSerTyrLeuAlaTh
3815 : CAATATGCCATTCAATGCTACGCTGGTGACTTCCACAATGAAGATCTCTCGTATCTTGCCAC : 3877

 748 : eGlyGluAspGlyPheSerAlaAsnAlaValArgThrGlnIleMetGluArgAsnIleGly :  767
        ||||||!  |||  !!!!:!!:!!||||||! !:!!:!!:!!   ..!! !|||  !|||
       rGlyGluAlaGlyThrThrSerHisAlaValIleSerLysValProArgLeuAsnGlyGly
3878 : AGGTGAAGCAGGTACCACCTCACACGCTGTCATTTCCAAAGTACCAAGGCTGAATGGTGGG : 3939
```
